# A modular mRNA–LNP vaccine platform enables integrated RNA, lipid and antigen design to protect against CCHFV

**DOI:** 10.64898/2026.01.17.699915

**Authors:** Touraj Farzani, Nallely Espinoza, Mahdiyeh M. Manafi, Stephen R. Welch, JoAnn D. Coleman-McCray, Virginia Aida-Ficken, Jessica R. Spengler, Éric Bergeron, Christina F. Spiropoulou, Clarice Borges, Kristine Bielecki, Lee Li, Devan Shah, Marietou Paye, Eugenia Rojas, Sophie N. Spector, Pedram Samani, Lisa E. Hensley, Al Ozonoff, Pardis C. Sabeti

## Abstract

mRNA–lipid nanoparticle (LNP) vaccines are programmable, multi-component systems in which immune outcomes emerge from coupled control of nanoparticle chemistry, RNA regulatory architecture, and antigen design. Here we establish an integrated engineering framework that quantitatively maps how ionizable lipid identity, untranslated region (UTR) configuration, and 5′ cap structure shape innate activation landscapes and thereby tune the magnitude, cellular distribution, and polarization of adaptive immunity. Benchmarking three ionizable lipids shows that lipid chemistry imprints distinct cytokine and chemokine milieus during dendritic cell–T cell priming that mirror downstream T cell activation phenotypes, identifying lipid structure as a determinant of pathway-selective activation; notably, our first-in-study lipid exhibits benchmark-comparable immunostimulatory profiles, supporting further translational evaluation. Using Crimean–Congo hemorrhagic fever virus (CCHFV) as a model high-consequence pathogen, we show that UTRs act as modular regulatory elements that redirect cytokine outputs, tuning effector versus proliferative programs and expanding helper polarization in an antigen-dependent manner. Cap structure functions primarily as a quantitative gain control, scaling cellular and humoral response magnitude without overriding antigen-defined polarization. Within this optimized platform, antigen architecture defines functional constraints on protection: structural modifications reshape immune hierarchies and antibody quality. Integrating these design axes yields an optimized mRNA–LNP vaccine encoding the CCHFV secreted glycoprotein complex (sGCs) that achieves 90% protection in an immunosuppressed murine lethal-challenge model with minimal clinical signs. Together, these data define generalizable design principles for rational, multi-parameter optimization of mRNA–LNP vaccines.

## Introduction

Emerging, pandemic-potential viruses require vaccine platforms that can be rapidly reconfigured while maintaining predictable performance across diverse antigens and deployment contexts (1). Crimean–Congo hemorrhagic fever virus (CCHFV), a tick-borne, high-consequence pathogen causing severe disease with unpredictable spread and no licensed vaccine, exemplifies this challenge (2). mRNA–lipid nanoparticle (LNP) vaccines offer an unprecedented platform for rapid vaccine design and deployment, enabling swift responses to emerging infectious threats (3). This speed is essential, but it is not sufficient: for many targets, mRNA vaccine potency, immune polarization and tolerability remain highly sensitive to variation in RNA regulatory architecture, antigen expression kinetics and nanoparticle chemistry (4). mRNA–LNP vaccines are multi-component systems whose performance emerges from interactions among RNA regulatory elements and nanoparticle chemistry, rather than from any single component alone (5). Within these systems, 5′ cap structures (6), untranslated regions (UTRs) (7) and nucleoside modifications (8) shape mRNA stability, translation, and innate sensing, while ionizable lipid composition governs cellular delivery, endosomal escape and inflammatory signaling (9).

Although individual mRNA-LNP vaccine components have been optimized in isolation, the field lacks an integrated framework that treats RNA and LNP design variables as coupled control knobs and quantitatively links their combinations to innate activation and adaptive immune fate decisions (10). As a result, generalizable design rules remain limited, and mRNA–LNP performance often varies unpredictably when extended beyond a narrow set of clinically validated targets. Current mRNA–LNP vaccine development relies on a narrow set of proprietary ionizable lipids (11), constraining independent optimization, manufacturing flexibility and systematic exploration of lipid chemistry (12). This reliance complicates efforts to disentangle how lipid structure contributes to innate immune activation and reactogenicity (13), particularly because accumulating evidence shows that LNP components can themselves stimulate innate pathways independent of the mRNA payload (9). Together, these considerations underscore the need for comparative, mechanistically grounded lipid benchmarking alongside RNA regulatory tuning.

Here we establish a modular molecular- and nanoscale-engineering framework for mRNA–LNP vaccines using CCHFV as a tractable model system (Figure 1). We systematically evaluate antigen architecture, including targeted structural modifications, mRNA regulatory design (cap and UTR configurations) and ionizable lipid chemistry in coordinated assays spanning expression, innate activation, cellular priming, humoral quality and in vivo efficacy. This integrated approach is designed to enable identification of a potent CCHFV vaccine candidate and to establish transferable design principles for rational, next-generation mRNA–LNP vaccine engineering.

**Figure 1.**
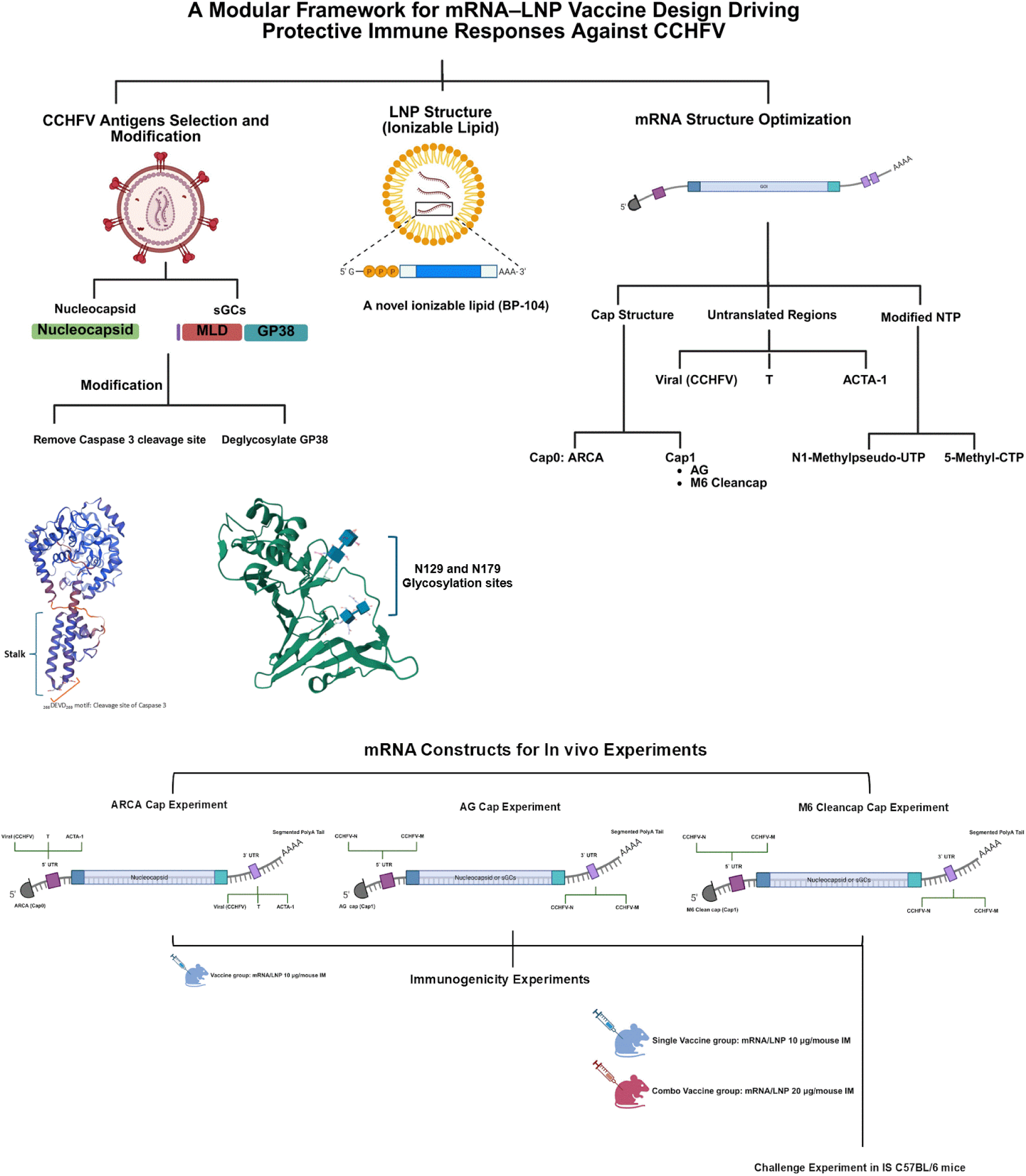
Modular mRNA–LNP design framework and in vivo construct set for CCHFV vaccination. **a,** Schematic overview of the integrated mRNA–LNP engineering framework spanning antigen architecture, ionizable lipid chemistry, and mRNA regulatory design. Two CCHFV immunogen classes were evaluated: the intracellular nucleocapsid (NP) and the secreted glycoprotein complex (sGCs; mucin-like domain (MLD)–GP38). Targeted antigen modifications included removal of a canonical caspase-3 cleavage motif in NP (NPmut) and deletion of two conserved GP38 N-linked glycosylation sites (GP38Δglyc; N129 and N179). LNP formulation incorporated the ionizable lipid BP-104. mRNA design variables included 5′ cap structure (Cap0 ARCA; Cap1 AG; Cap1 CleanCap M6), untranslated regions (UTRs; viral/CCHFV, T, and ACTA-1), and modified nucleotides (N1-methylpseudouridine-UTP and 5-methyl-CTP). Ionizable lipid chemistry sets the innate cytokine/chemokine context, UTR architecture tunes T cell cytokine polarization and cellular distribution, cap structure scales response magnitude and antibody output, and antigen architecture defines the protective ceiling. **b,** mRNA construct configurations used for in vivo studies, grouped by cap-structure experiments. ARCA-capped constructs were paired with alternative UTRs (viral/CCHFV, T, ACTA-1), whereas AG- and CleanCap M6–capped constructs used defined 5′/3′ UTR configurations (CCHFV-N and CCHFV-M) and a segmented poly(A) tail. Immunogenicity studies used intramuscular vaccination with 10 µg mRNA/LNP per mouse for single-antigen groups and 20 µg total mRNA/LNP for combination groups. Protective efficacy was evaluated in transiently immunosuppressed C57BL/6 mice in a lethal CCHFV challenge model.

## Results

### Antigen selection and modular mRNA design space

To compare intracellular and secreted viral proteins within the mRNA–LNP platform, we selected Crimean–Congo hemorrhagic fever virus (CCHFV) antigens spanning distinct cellular localizations. As a representative intracellular antigen, we focused on the nucleocapsid protein (NP), which is highly immunogenic and consistently elicits strong T cell responses across vaccine platforms (14–16). As a representative surface antigen, we included the secreted glycoprotein complex (sGCs; mucin-like domain–GP38) (17), given the central role of glycoprotein-directed immunity in antibody-mediated protection for many viral pathogens (18). We then introduced targeted antigen variants to test how defined structural perturbations reshape immune priming. For NP, we removed a canonical caspase-3 cleavage site (19) to generate NPmut, enabling evaluation of altered intracellular processing (20, 21). For sGCs, we deleted two conserved N-linked glycosylation sites on GP38 to generate sGCs(GP38Δglyc), allowing assessment of epitope accessibility (22). The antigen set evaluated throughout the study therefore consisted of NP, NPmut, sGCs, and sGCs(GP38Δglyc).

In parallel, to define how mRNA regulatory architecture shapes immune outcomes, we systematically varied key transcript features while holding other parameters constant. We compared 5′ cap structures (Cap0: ARCA (Supplementary Figure 3a); Cap1: AG (Supplementary Figure 3b) and CleanCap M6 (Supplementary Figure 3c)), and confirmed antigen expression from M6 Cleancapped mRNAs by immunoblot (Supplementary Figure 3d). We further evaluated UTR configurations (CCHFV-derived, T, and ACTA-1) and dual nucleoside modifications (N1-methylpseudouridine and 5-methylcytidine), while maintaining a uniform 110-bp poly(A) tail. Together, these variables defined an mRNA design axis that, combined with the antigen panel, enabled integrated assessment of how transcript architecture and antigen configuration influence expression, innate activation, and adaptive immune responses.

### Physicochemical benchmarking identifies BP-104 as a potential formulation lipid

Because ionizable lipid chemistry is a key determinant of LNP performance (23), we compared SM-102 and ALC-0315 with BP-104 (Supplementary Figure 4a), a newly introduced ethanolamine-headgroup lipid structurally related to SM-102. Across constructs, BP-104 formed stable, monodisperse nanoparticles with polydispersity indexes below 0.15 (Supplementary Figure 4c), near-neutral zeta potentials (–8 ± 1 mV), and ∼90% mRNA encapsulation efficiency. BP-104 LNPs exhibited apparent pKa values of 6.6 to 6.9 (Supplementary Figure 4d), consistent with efficient endosomal escape and strong immunogenicity (24). Empty BP-104 particles caused minimal cytotoxicity (Supplementary Figure 3e), whereas NP/LNP showed slightly higher effects than NPmut/LNP, indicating that mRNA sequence and lipid chemistry can jointly influence tolerability. Consistent with these in vitro observations, no overt histopathological abnormalities were observed in representative liver sections at endpoint, and cleaved caspase-3 staining was low/rare across AG-capped groups (Supplementary Figure 6). Collectively, these results show that BP-104 supports well-behaved nanoparticle formation (Supplementary Figure 4b) with favorable physicochemical and safety properties for vaccine formulation.

### BP-104 elicits benchmark-comparable innate pathway activation

To benchmark innate activation (25, 26), we encapsulated NPmut mRNA in LNPs containing BP-104, SM-102, or ALC-0315 and introduced these formulations into HEK293/Blue and THP1-Dual reporter cells. All three lipids induced robust type I interferon (Supplementary Figure 5c) and IL-6 (Supplementary Figure 5b) signaling after 24 h, with ALC-0315 producing slightly higher IFN-I responses at some doses. Empty LNPs also triggered signaling, indicating contributions from both lipid and mRNA to innate stimulation. NF-κB activation (Supplementary Figure 5a) mirrored IFN-I and IL-6 patterns, and BP-104 elicited responses comparable to the two clinically used lipids. These data indicate that BP-104 engages canonical innate pathways at effective, non-excessive levels, consistent with balanced potency and tolerability.

### UTR architecture differentially tunes DC-driven CD4⁺ and CD8⁺ priming in an antigen-dependent manner

Given the physicochemical and innate benchmarking supporting BP-104 as a formulation platform, we next examined how transcript-level regulatory design modulates adaptive priming. BP-104 mRNA–LNPs encoding NP were delivered to murine dendritic cells (DC2.4 cells), co-cultured with naïve CD4⁺ or CD8⁺ T cells, and intracellular cytokine responses were quantified (Figure 2). For NP, most UTR configurations produced relatively uniform CD8⁺ IFN-γ and TNF-α levels (Figure 2b), indicating limited sensitivity of cross-priming to transcript context. However, specific UTRs emerged as outliers that enhanced IFN-γ/TNF-α or IL-2, consistent with tunable effector versus proliferative programs. In contrast, NP-driven CD4⁺ responses (Figure 2a) showed greater variability, with several UTRs inducing concurrent IL-4 and IL-17A alongside Th1 cytokines, demonstrating that regulatory elements can broaden helper polarization even for an intracellular antigen.

**Figure 2.**
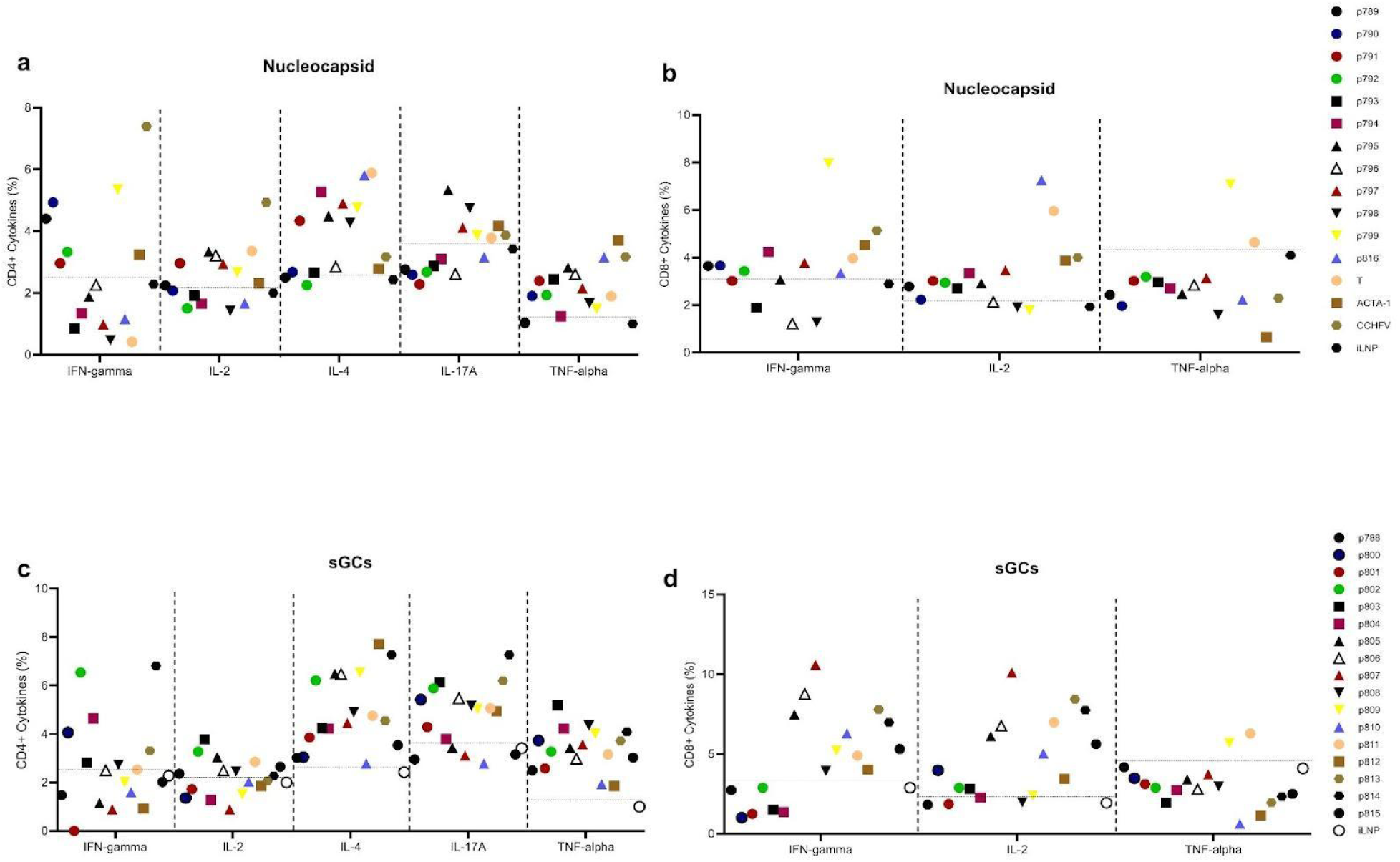
Untranslated region (UTR) architecture differentially programs cytokine polarization in CD4⁺ and CD8⁺ T cells in an antigen-dependent manner. Murine dendritic cells (DCs) were transfected with BP-104 mRNA/LNPs encoding either nucleocapsid (NP; *a,b*) or the secreted glycoprotein complex (sGCs; *c,d*) containing distinct 5′ and 3′ untranslated regions (UTRs), then co-cultured with naïve CD4⁺ (*a,c*) or CD8⁺ (*b,d*) T cells. Intracellular cytokine staining was performed after stimulation to quantify the frequency of cytokine-positive T cells (IFN-γ, IL-2, IL-4, IL-17A, and TNF-α). Each symbol represents a unique UTR variant, as indicated in the key. Horizontal dotted lines denote the mean baseline response from iLNP (empty LNP) controls. For NP mRNAs (*a,b*), CD8⁺ T cell responses were relatively uniform across most UTRs, with select variants enhancing IFN-γ/TNF-α or IL-2 production, whereas CD4⁺ T cells exhibited broader cytokine diversification, including induction of IL-4 and IL-17A. For sGC mRNAs (*c,d*), UTR identity strongly influenced both magnitude and composition of responses. Multiple constructs significantly increased CD8⁺ IFN-γ and IL-2 frequencies above iLNP baseline, while CD4⁺ responses showed concurrent Th1-, Th2-, and Th17-associated cytokine profiles. Together, these results demonstrate that UTR regulatory architecture tunes both the amplitude and polarization of T cell priming in an antigen-specific manner, with intracellular (NP) and secreted (sGCs) antigens exhibiting distinct UTR-sensitivity landscapes.

We next evaluated UTR effects on a secreted antigen using sGCs mRNA–LNPs in the same assay. In this context, UTR configuration exerted a stronger influence on CD8⁺ priming (Figure 2d), with certain UTRs substantially increasing IFN-γ and IL-2 above baseline, while TNF-α varied less across contexts.CD4⁺ responses (Figure 2c) displayed mixed Th1/Th17/Th2 profiles, indicating that for secreted glycoproteins, UTRs modulate both response magnitude and helper diversity. Together, these results identify UTR engineering as a practical lever for tuning DC-driven priming in an antigen-specific manner.

### Ionizable lipid identity shapes the inflammatory milieu and Th1/Th17 polarization during antigen presentation

We next asked whether ionizable lipid composition similarly influences antigen presentation and T cell polarization (27). Primary murine DCs were loaded with NPmut mRNA formulated in BP-104, SM-102, or ALC-0315 LNPs and co-cultured with naïve T cells. All formulations promoted Th1/Th17-skewed activation (Figure 3), with strong IFN-γ and TNF-α production in both CD4⁺ and CD8⁺ compartments (Figure 3a, b) and the emergence of a distinct IL-17A⁺ CD4⁺ subset (Figure 3d); negative controls remained minimal. Among lipids, ALC-0315 induced the highest frequencies of IFN-γ⁺ and TNF-α⁺ T cells and the strongest IL-17A⁺ CD4⁺ responses, whereas BP-104 elicited intermediate activation and SM-102 produced the weakest responses. IL-2⁺ CD8⁺ T cell frequencies (Figure 3c) also varied significantly by lipid, consistent with differential proliferative programming.

**Figure 3.**
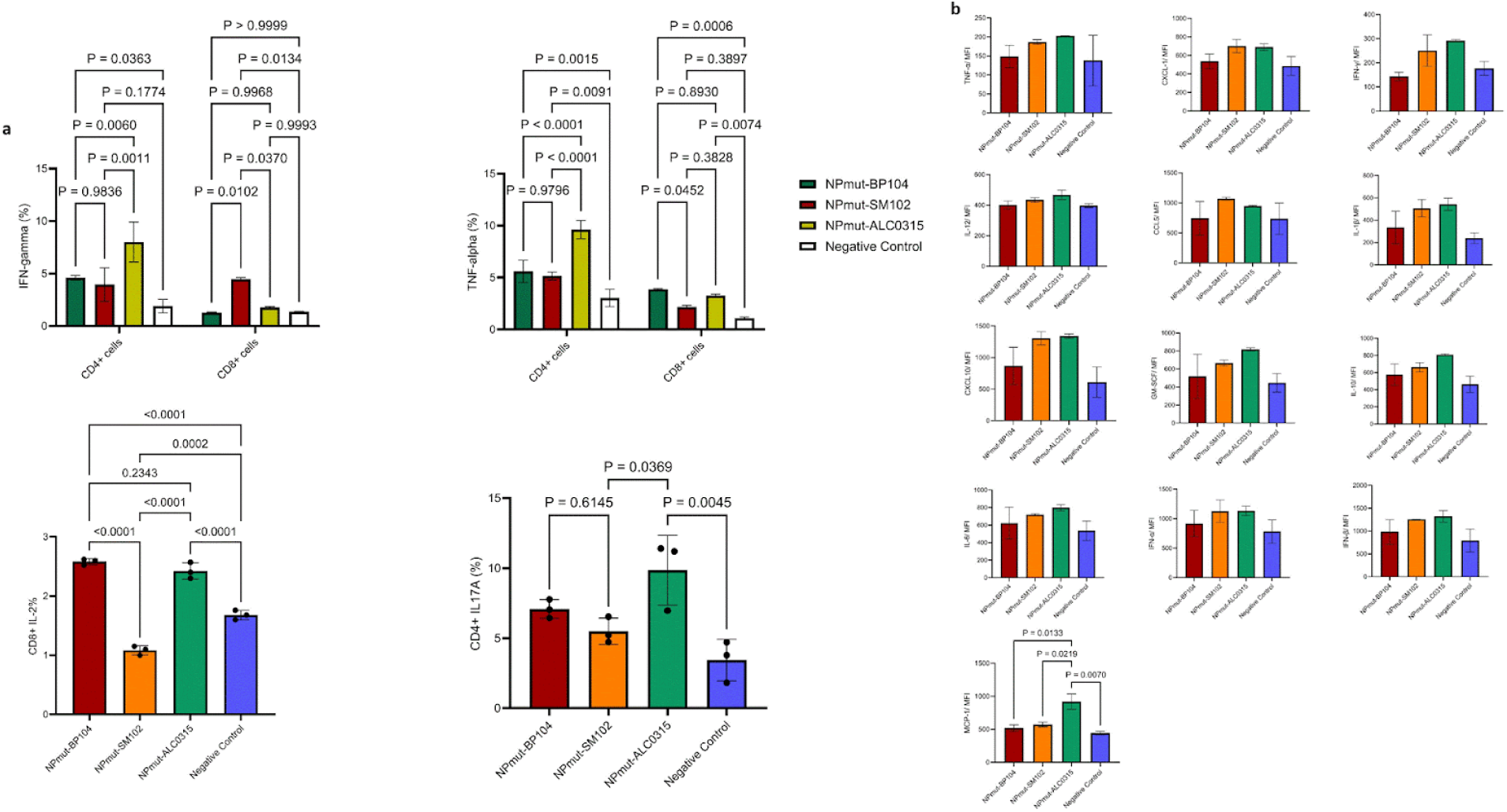
Ionizable lipid chemistry tunes dendritic cell–mediated antigen presentation and downstream T cell polarization. Primary murine dendritic cells (DCs) were loaded with NPmut mRNA formulated in lipid nanoparticles (LNPs) containing BP-104, SM-102, or ALC-0315, then co-cultured with naïve CD4⁺ and CD8⁺ T cells to quantify intracellular cytokines and soluble mediators in supernatants. **a,** Frequencies of cytokine-positive CD4⁺ and CD8⁺ T cells measured by intracellular cytokine staining, including IFN-γ and TNF-α (top), IL-2⁺ CD8⁺ cells (bottom left), and IL-17A⁺ CD4⁺ cells (bottom right). Bars show group means with individual biological replicates overlaid; statistical comparisons and corresponding *P* values are indicated. Across formulations, NPmut mRNA/LNP-loaded DCs elicited Th1-skewed responses (IFN-γ and TNF-α) with a measurable IL-17A⁺ CD4⁺ population, whereas negative control cultures remained low. ALC-0315 drove the strongest IFN-γ⁺ and TNF-α⁺ T cell responses across CD4⁺ and CD8⁺ compartments and increased IL-17A⁺ CD4⁺ frequencies, BP-104 produced intermediate activation, and SM-102 was comparatively attenuated. Lipid-dependent differences were also evident for IL-2⁺ CD8⁺ responses, indicating differential programming of proliferative/expansion-associated outputs. **b,** Secreted cytokine and chemokine profiles in DC–T cell co-culture supernatants (mean fluorescence intensity, MFI) measured for inflammatory and recruitment-associated mediators, including TNF-α, CXCL1, IFN-γ, IL-12, CCL5, IL-4, CXCL10, GM-CSF, IL-10, IL-6, IP-10, and MCP-1. Error bars indicate variability across replicates; selected pairwise comparisons are annotated with *P* values. Supernatant signatures mirrored intracellular T cell phenotypes: ALC-0315 induced the most pronounced inflammatory/recruitment program, BP-104 generated a coherent but moderated profile, similar to SM-102. Together, these data show that ionizable lipid identity imprints innate cytokine/chemokine cues that scale and shape downstream Th1/Th17 polarization under fixed mRNA payload and DC-loading conditions.

To define how lipid chemistry shapes the inflammatory context during antigen presentation (28), we quantified cytokine and chemokine profiles produced by DCs under the same conditions (Figure 3e). ALC-0315 produced the most inflammatory profile, including elevated IL-12, IL-6, CXCL10, and MCP-1, whereas BP-104 generated moderate, coordinated responses and SM-102 remained low across analytes. Collectively, these data establish lipid chemistry as a decisive variable controlling both delivery efficiency and the inflammatory milieu that instructs Th1/Th17 polarization. Under identical payload conditions, ALC-0315 maximized activation, BP-104 balanced potency and inflammation, and SM-102 produced the mildest stimulation (Figure 3).

### In vivo, UTR architecture programs NP-specific cellular immunity under ARCA capping

To connect mRNA regulatory features to in vivo immunity, we immunized mice with BP-104 mRNA–LNPs varying in cap and UTR configuration. Among ARCA-capped constructs (Figure 4a), transcript architecture strongly influenced both response magnitude and phenotype when stimulated with correspondent antigen (Supplementary Figure 1a, b). NP constructs bearing the T-UTR favored CD8⁺ IFN-γ, IL-2, and TNF-α responses (Figure 4c), whereas the ACTA-1 UTR promoted robust CD4⁺ Th1/Th17 cytokine production (Figure 4d). ELISPOT assays confirmed broad IL-2 responses and the highest IFN-γ levels for ACTA-1 and CCHFV-derived UTRs (Figure 4b), demonstrating that UTR design (Supplementary Figure 2a) is a powerful determinant of cellular polarization in vivo.

**Figure 4.**
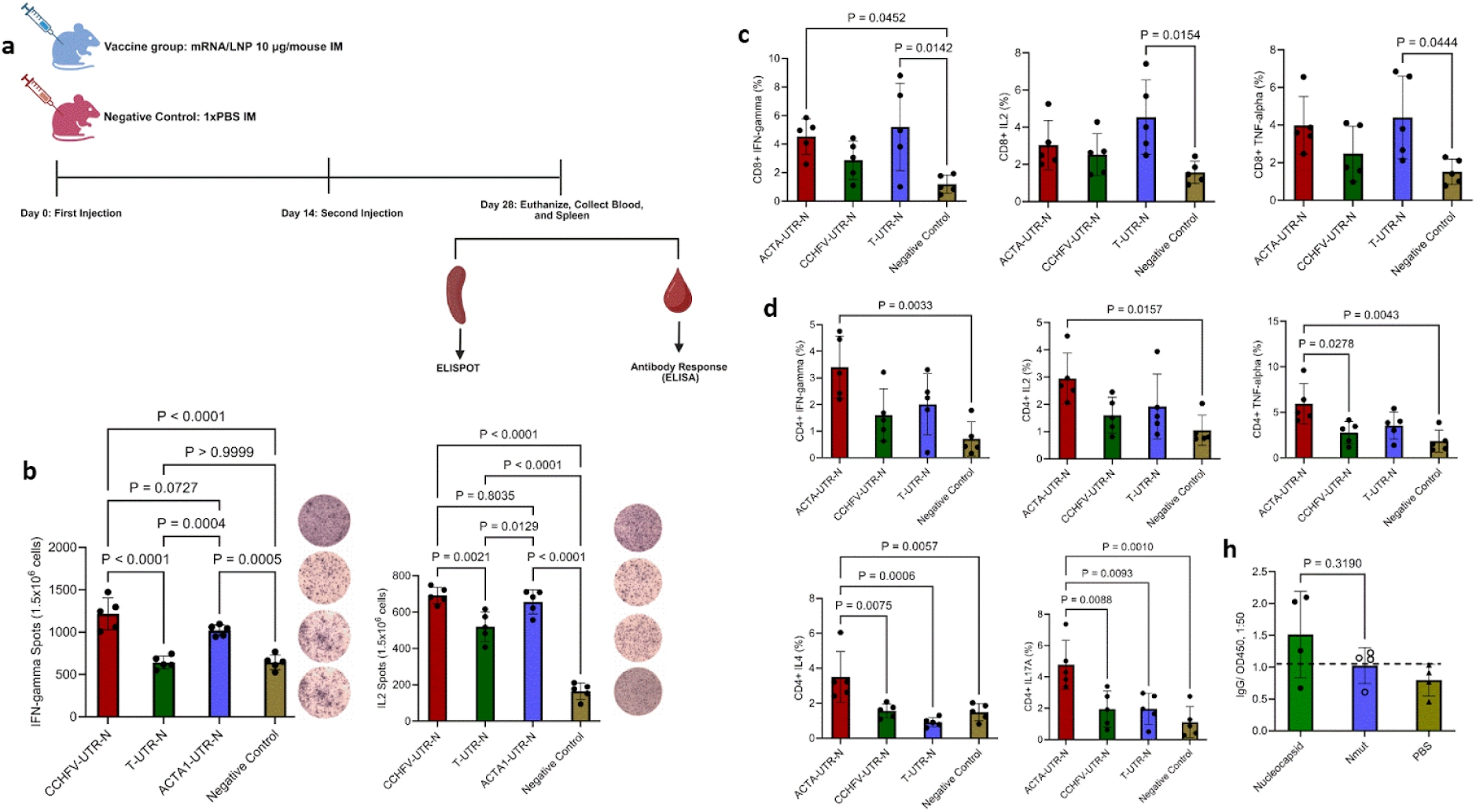
UTR architecture modulates NP-specific cellular immunity in vivo under ARCA capping. **a,** Immunization and sampling scheme. Female C57BL/6J mice (8–10 weeks old; n = 5 per group) were immunized intramuscularly with 10 μg of ARCA-capped nucleocapsid (NP) mRNA–LNP on days 0 and 14 using three UTR configurations (CCHFV-UTR–NP, T-UTR–NP, or ACTA-1–UTR–NP). Negative-control animals received 1× PBS intramuscularly. On day 28, animals were euthanized and blood and spleens were collected for serology (ELISA) and cellular immunogenicity assays (ELISPOT and intracellular cytokine staining). **b,** Antigen-specific cytokine secretion by ELISPOT. Splenocytes harvested on day 28 were stimulated for 24 h with recombinant CCHFV NP (10 μg ml⁻¹) and analyzed for IFN-γ and IL-2 spot-forming cells (SFCs; normalized per 1.5 × 10⁶ input cells). Representative ELISPOT wells are shown adjacent to quantification. All NP mRNA–LNP vaccines induced significantly higher IL-2 secretion than stimulated negative controls. Constructs containing CCHFV and ACTA-1 UTRs elicited significantly greater IL-2 responses than the T-UTR construct and negative controls, whereas IFN-γ responses were also significantly increased relative to controls, with UTR-dependent differences indicated by the displayed P values. **c,** CD8⁺ T cell cytokine production. Day-28 splenocytes were stimulated for 5 h with recombinant NP (10 μg ml⁻¹) and analyzed by intracellular cytokine staining (ICS) to quantify frequencies of IFN-γ⁺, IL-2⁺, and TNF-α⁺ CD8⁺ T cells. The T-UTR construct increased CD8⁺ IFN-γ, IL-2, and TNF-α relative to negative controls, although the three vaccine candidates did not differ significantly from one another across these CD8⁺ readouts (as indicated by the annotated P values). **d,** CD4⁺ T cell cytokine polarization. Day-28 splenocytes were stimulated for 5 h with recombinant NP (10 μg ml⁻¹) and analyzed by ICS to quantify IFN-γ⁺, IL-2⁺, IL-4⁺, TNF-α⁺, and IL-17A⁺ CD4⁺ T cells. The ACTA-1–UTR construct induced markedly higher frequencies of all five cytokines compared with the CCHFV-UTR and T-UTR constructs and the negative-control group, indicating broadened helper polarization. CCHFV-UTR– and T-UTR–based constructs produced comparable but lower cytokine frequencies in CD4⁺ T cells. Bars represent group means with individual animals overlaid as points; error bars denote variability within groups. Statistical analyses were performed in GraphPad. Between-group comparisons used one-way ANOVA with Tukey’s multiple comparisons test; when normality assumptions were not met, the Kruskal–Wallis test was used. A significance threshold of P < 0.05 was applied unless otherwise indicated, and exact P values are shown in the figure.

### Cap structure scales response magnitude without overriding antigen-defined polarization

Having established that UTR/transcript architecture can reprogram response magnitude and cellular distribution in vivo, we compared AG- and CleanCap M6–capped constructs across antigens using CCHFV-derived UTRs (Supplementary Figure 2b) to separate regulatory effects from antigen-driven bias.

Under AG capping (Figure 5a), NP and NPmut induced Th1-biased ELISPOT IFN-γ responses, whereas sGCs and sGCs(GP38Δglyc) produced ELISPOT IL-4–dominant Th2 responses (Figure 5b), indicating that antigen identity dictates qualitative polarization under matched regulatory context. CleanCap M6 constructs (Figure 6a) further increased total T cell responses without altering antigen-specific bias: M6-capped NP and NPmut elevated CD8⁺ IFN-γ, while M6-capped sGCs(GP38Δglyc) enhanced CD4⁺ IL-2 and TNF-α production (Figure 6c). IL-4 remained undetectable across conditions, and ELISPOT assays confirmed strong IFN-γ induction by NP antigens (Figure 6b). Together, these findings indicate that cap and UTR architecture modulate response amplitude and cellular distribution, whereas antigen structure defines polarization identity.

**Figure 5.**
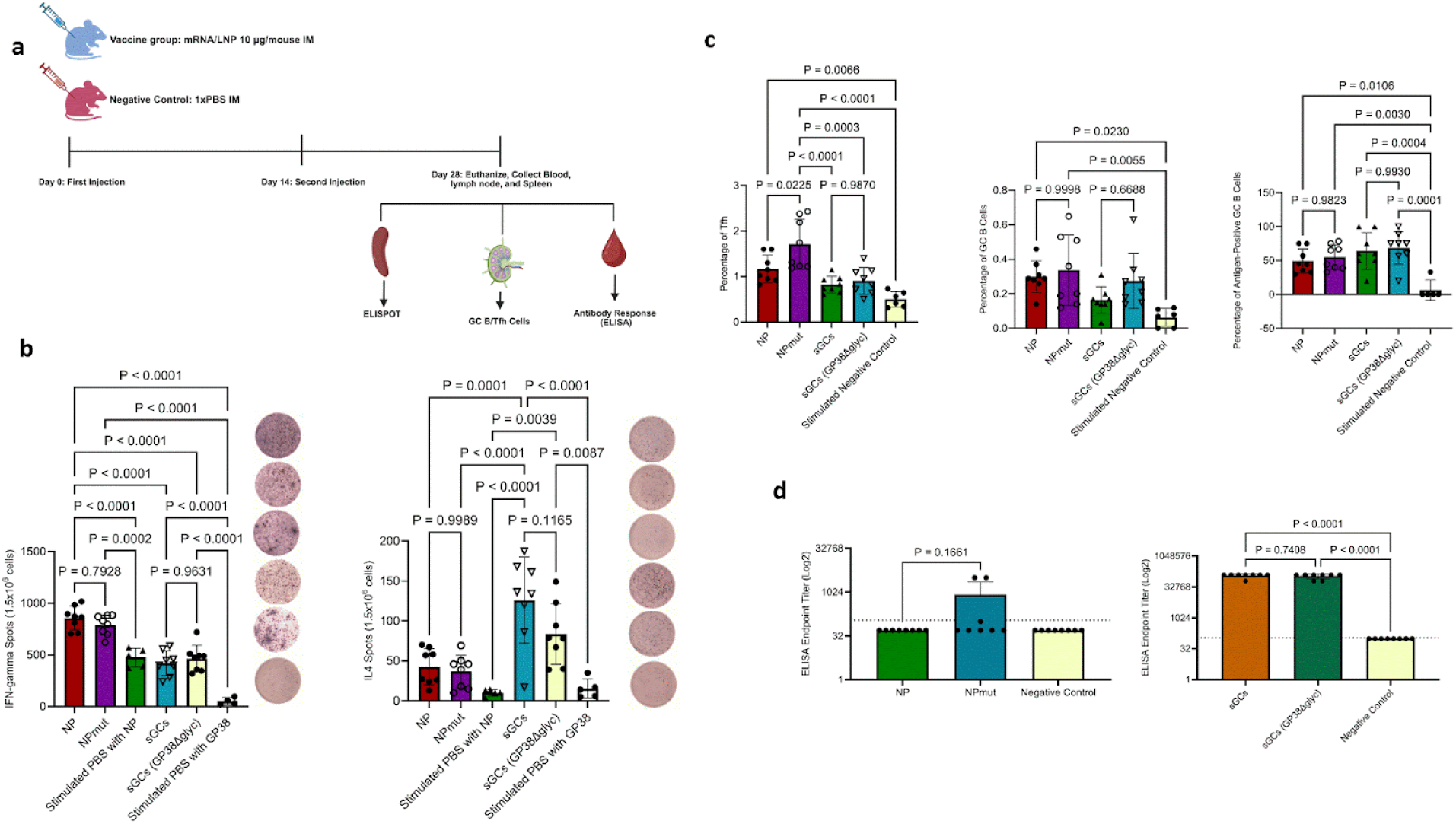
Immunogenicity of AG capped nucleocapsid, NPmut, sGCs and sGCs (GP38Δglyc) mRNAs with CCHFV UTR region in C57BL/6J mice model in a booster regime. **a,** The immunization schedule and experimental workflow are shown for AG-capped mRNA–LNP formulations encoding nucleocapsid (NP), mutant nucleocapsid (NPmut), sGCs, and sGCs (GP38Δglyc). Female C57BL/6J mice (8–10 weeks old) were immunized intramuscularly with 10 µg of each mRNA–LNP vaccine on days 0 and 14. Blood, spleens, and inguinal lymph nodes were collected on day 28 for downstream immunological analyses, including ELISA, germinal center (GC) B and T follicular helper (Tfh) cell profiling, and ELISPOT assays. **b,** Splenocytes were stimulated for 24 hours with 10 µg/mL of recombinant nucleocapsid or GP38 proteins and analyzed by ELISPOT to quantify IFN-γ and IL-4 secretion. The NP and NPmut groups exhibited significantly higher IFN-γ–secreting cell frequencies compared with sGCs, sGCs (GP38Δglyc), and negative control groups, whereas sGCs and sGCs (GP38Δglyc) formulations elicited stronger IL-4 responses than NP, NPmut, and controls. **c,** GC B and Tfh cell populations were quantified in lymph nodes collected on day 28. The NPmut group displayed a significantly increased percentage of GC B and Tfh cells relative to negative controls, while the NP group showed a higher frequency of GC B cells compared with controls. Antigen-specific GC B cells were elevated across all vaccine groups relative to controls, with no significant differences among vaccine formulations. **d,** Antibody levels measured by in-house ELISA on day 28 showed that two out of eight mice in the NPmut group had detectable antigen-specific antibodies, while all samples in the sGCs and sGCs (GP38Δglyc) groups exhibited significantly higher antibody titers compared with negative controls. No significant differences were observed between sGCs and sGCs (GP38Δglyc) groups. Statistical analysis was performed using one-way ANOVA followed by Tukey’s post-hoc multiple comparisons test. Significant differences are shown as *p*-values.

**Figure 6.**
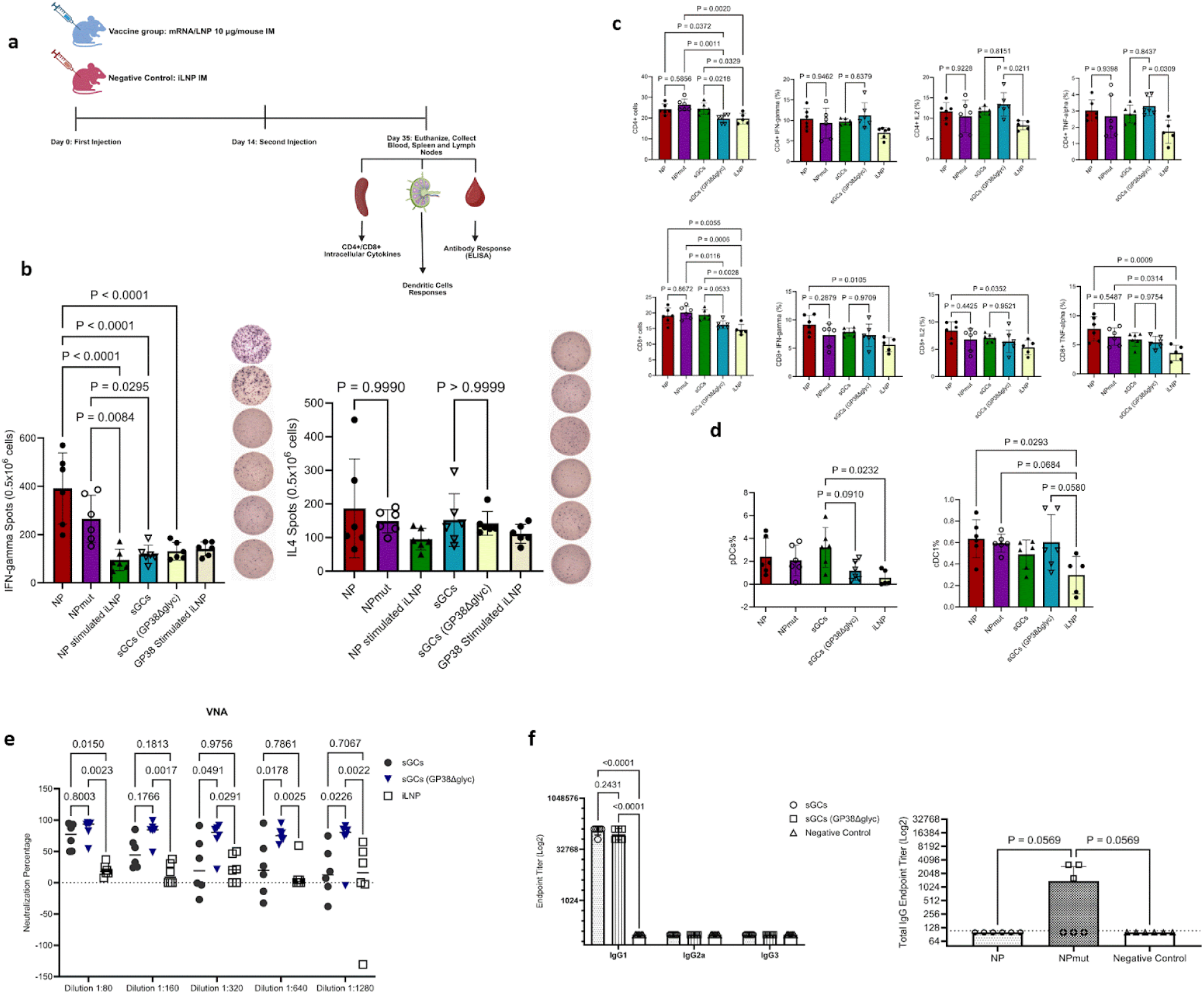
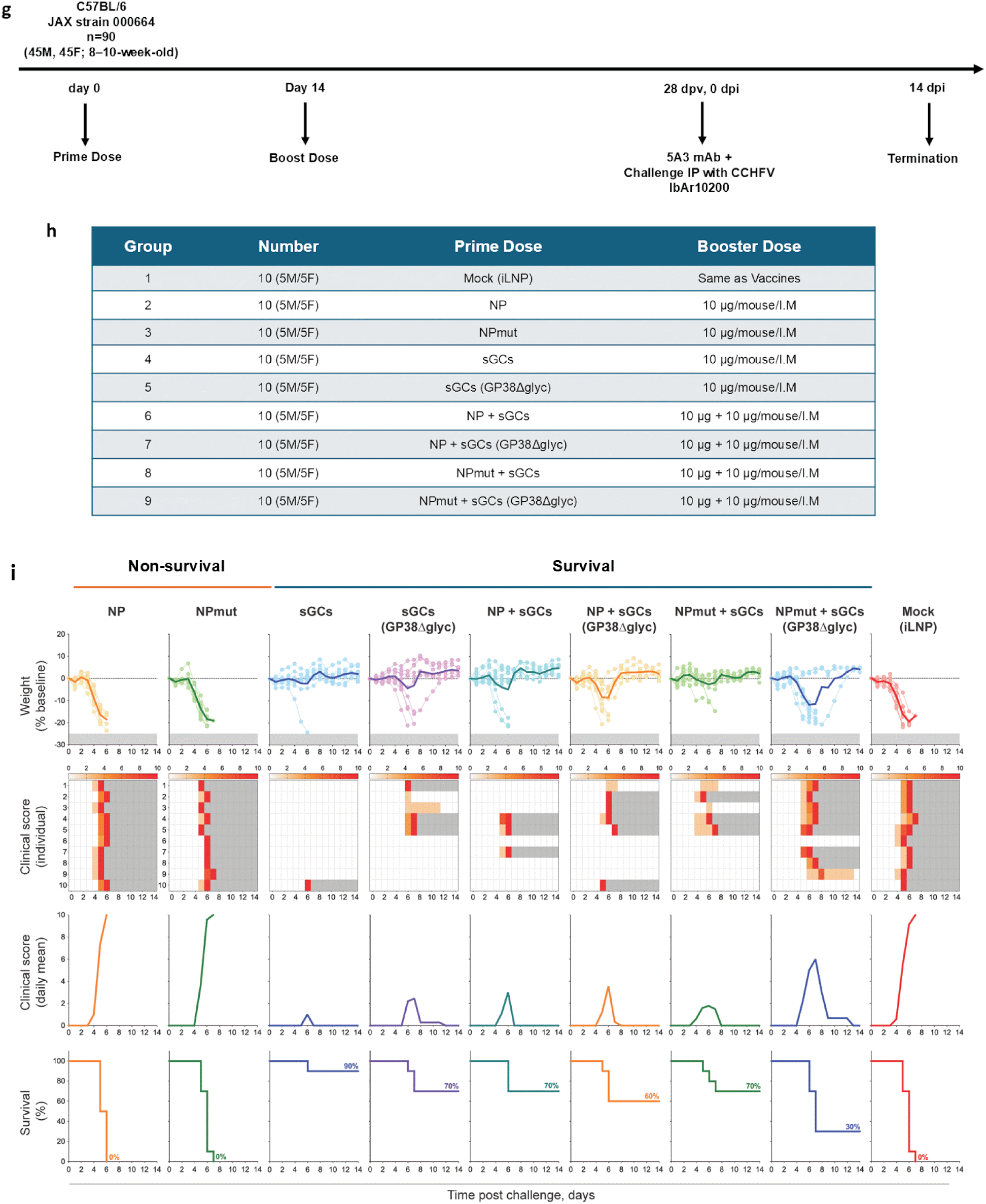
Immunogenicity and protection efficacy of Cleancap M6 mRNA–LNP vaccines encoding CCHFV antigens. **a,** Female C57BL/6J mice were immunized intramuscularly with Cleancap M6 mRNA–LNP formulations encoding nucleocapsid (NP), mutant nucleocapsid (NPmut), sGCs, or sGCs (GP38Δglyc). Blood, spleens, and lymph nodes were collected on day 35 for downstream analyses, including ELISA, dendritic cell profiling, and ELISPOT assays. **b,** Splenocytes were stimulated for 24 hours with 10 µg/mL of nucleocapsid or GP38 proteins and analyzed by ELISPOT for IFN-γ and IL-4 secretion. NP and NPmut groups induced significantly higher IFN-γ responses compared with sGCs, sGCs (GP38Δglyc), and negative control, whereas sGCs and sGCs (GP38Δglyc) showed no significant IFN-γ response. None of the vaccine groups elicited significant IL-4 production. **c,** Splenocytes were stimulated ex vivo for 24 hours with 10 µg/mL of nucleocapsid or GP38 proteins and analyzed for intracellular cytokines IFN-γ, IL-2, and TNF-α in CD8⁺ and CD4⁺ T cells. All vaccine groups elicited robust CD4⁺ and CD8⁺ T cell responses compared with negative controls, except for sGCs (GP38Δglyc). NP vaccination induced significantly higher cytokine production in CD8⁺ T cells relative to control, whereas sGCs (GP38Δglyc) uniquely enhanced IL-2 and TNF-α in CD4⁺ T cells. No significant differences were observed among vaccine groups. **d,** Lymph nodes were analyzed for plasmacytoid dendritic cells (pDCs) and conventional CD11c⁺ DCs (cDC1). Only sGCs and NP vaccine groups demonstrated significantly increased pDC and cDC1 frequencies, respectively, relative to controls. **e,** Dots represent the distribution of individual serum samples at each dilution. Both immunogens elicited robust neutralizing activity at low serum dilutions (1:100–1:200). At higher dilutions (1:1600–1:6400), sera from mice immunized with MLD-GP38Δglyc exhibited significantly greater neutralization than the wild-type MLD-GP38 group, indicating that removal of GP38 glycans enhances the potency and/or durability of the neutralizing antibody response. Negative control sera (blue) showed minimal inhibition across all dilutions. **f,** ELISA revealed that 3 out of 6 mice in the NPmut group were positive, while no NP-vaccinated mice showed detectable antibody. Both sGCs and sGCs (GP38Δglyc) groups exhibited significantly higher antibody responses than negative controls, with no differences between these two groups. **g and h,** The immunization/grouping scheme in IS C57BL/6 mice. **i,** Weight, clinical score and survival of C57BL/6J mice vaccinated with CCHFV based mRNAs 35 days prior to challenge and provided a boost dose at 21 days prior to challenge (n = 10 per group). On the day of challenge, mice were transiently immunosuppressed with anti-mouse IFNAR-1 mAb IP and subsequently inoculated with CCHFV strain IbAr10200 SC. Weights depicted as percent change from baseline at D0. Clinical scores are shown as individual daily scores (0–10; increasing intensity of red indicates higher scores, and gray boxes denote end of monitoring due to euthanasia or reaching a lethal endpoint; top) or as mean scores for each experimental group (bottom).

### Antigen architecture differentially governs Tfh and germinal center recruitment

Because cap/UTR and antigen-dependent T cell programs can propagate into humoral help, we profiled draining lymph nodes to quantify Tfh and germinal center (GC) responses (29). Draining lymph nodes were analyzed two weeks post-boost from mice immunized with AG-capped mRNA–LNPs encoding NP, NPmut, sGCs, or sGCs(GP38Δglyc). All vaccines significantly expanded Tfh cells relative to controls, with NPmut inducing the highest frequencies, exceeding NP and both sGC constructs. Despite this preferential Tfh expansion by NP-based antigens, antibody responses remained modest. In contrast, sGCs(GP38Δglyc) elicited the strongest antigen-specific GC B-cell responses, surpassing NP, NPmut, and sGCs, although total GC frequencies were similar across groups (Figure 5c). These results indicate that antigen architecture can independently govern Tfh versus GC recruitment, with NP favoring Tfh expansion and sGCs(GP38Δglyc) enhancing antigen-specific GC formation.

### Cap structure increases antibody magnitude while preserving antigen-imposed class bias; GP38 glycan removal enhances neutralization potency

To assess how mRNA cap structure influences antibody production, we measured serum IgG responses following ARCA-, AG-, or CleanCap M6–capped vaccinations by ELISA and pseudovirus neutralization. ARCA-capped constructs elicited weak antibody responses with pronounced strain dependence: C57BL/6J mice showed no detectable anti-NP IgG, whereas BALB/c mice exhibited low NP and NPmut titers at a 1:50 dilution (Figure 4h). Under AG capping, antibody responses tracked germinal center trends: NP remained antibody-negative, NPmut produced low responses, and both sGCs constructs generated strong total IgG responses, aligning antibody magnitude with GC recruitment rather than Tfh abundance (Figure 5d). CleanCap M6 further increased antibody titers across all antigens while preserving antigen-specific differences. NPmut modestly exceeded NP without statistical significance, whereas sGCs and sGCs(GP38Δglyc) induced dominant IgG1 responses that were significantly higher than controls and comparable between variants. IgG2a and IgG3 remained near baseline across conditions, indicating a polarized IgG1 profile (Figure 6f). These observations support a model in which cap structure scales antibody magnitude but does not override antigen-defined constraints on humoral class or strength.

Functional neutralization assays using vesicular stomatitis virus pseudotypes expressing truncated CCHFV glycoprotein (30) showed that sera from sGC-immunized mice neutralized virus at dilutions exceeding 1:160, a protective benchmark established previously (18). Neutralization persisted across serial dilutions and was consistently stronger for sGCs(GP38Δglyc), indicating that glycan deletion enhances exposure of neutralizing epitopes. Sera from mock iLNP groups failed to reach protective thresholds, confirming antigen dependence (Figure 6e). Collectively, these data show that sGC-encoding mRNA–LNPs elicit potent, functional antibodies meeting operational protection criteria (18), and that GP38 glycan removal increases neutralizing potency without compromising IgG1 dominance.

### Antigen-dependent shifts in draining lymph node DC subsets under CleanCap M6 vaccination

Because innate immune composition in draining lymph nodes can influence downstream immunity (31), we analyzed DC subsets following CleanCap M6 vaccination, the regimen selected for challenge studies. Antigen identity shaped DC distribution: sGCs increased plasmacytoid DC (pDC) frequencies, whereas NP vaccination preferentially expanded conventional type 1 DC (cDC1) populations (Fig. 6d). These antigen-specific shifts indicate that early innate environments differ across antigens even under identical mRNA–LNP delivery conditions.

### Protective efficacy in lethal challenge aligns with sGCs-driven humoral immunity

Finally, we tested eight CleanCap M6–capped mRNA–LNP regimens in a lethal CCHFV challenge model to link immunogenicity with protection (Figure 6g). C57BL/6J mice (n = 10 per group) (Figure 6h) were immunized intramuscularly with prime and boost doses, transiently immunosuppressed with anti-IFNAR1 antibody (MAR1-5A3), and challenged with the IbAr10200 strain. We compared four single-antigen vaccines (NP, NPmut, sGCs, sGCs(GP38Δglyc)) and four combination regimens (NP+sGCs, NP+sGCs(GP38Δglyc), NPmut+sGCs, NPmut+sGCs(GP38Δglyc)) with empty-LNP (iLNP) control, and monitored animals for 14 days post-challenge. Vaccination with sGCs alone conferred robust protection, with 90% survival and minimal clinical signs, whereas sGCs(GP38Δglyc) achieved 70% survival. In contrast, NP and NPmut conferred no protection, paralleling mock controls, all of which succumbed within 5–7 days. Combination regimens yielded intermediate, non-additive protection (30–70%), and none surpassed sGCs alone. Surviving animals were asymptomatic or displayed only mild, transient illness (Figure 6i). Together, these results show that under CleanCap M6 conditions, sGCs-based immunity is sufficient for protection, whereas NP-driven cellular responses are not, and that protective efficacy aligns with antibody quality and germinal center recruitment rather than T cell magnitude in this model.

## Discussion

Our study frames mRNA–LNP vaccines as a quantitatively navigable, multi-parameter design space in which immune outputs are emergent properties of coupled control knobs—nanoparticle chemistry, RNA regulatory architecture, and antigen configuration—rather than consequences of any single component in isolation. By applying a structured engineering strategy in a CCHFV model, we systematically perturbed these axes to map how lipid-imprinted innate cues and RNA-encoded translation control jointly shape response magnitude, cellular distribution, polarization, and protection, establishing a platform-forward blueprint for rational optimization of potency, inflammatory burden, and efficacy.

Within this platform, ionizable lipid chemistry emerged as a high-leverage, independent determinant of pathway-selective activation that propagates from innate signaling into adaptive fate decisions (32, 33). Despite comparable delivery efficiency, ALC-0315, SM-102, and the first-use BP-104 lipid imprinted distinct cytokine and chemokine landscapes during DC–T cell priming: ALC-0315 drove the strongest inflammatory/recruitment program and Th1/Th17 skewing, SM-102 was attenuated, and BP-104 occupied an intermediate regime characterized by coherent priming with moderated inflammatory output. Importantly, BP-104 engaged canonical innate pathways at levels comparable to clinically used lipids while supporting well-behaved nanoparticle properties, illustrating that delivery efficiency and inflammatory burden can be decoupled and expanding the formulation/manufacturing space beyond a narrow set of proprietary lipids.

RNA regulatory architecture provided a second, orthogonal set of tuning parameters. UTR selection programmably reweighted cytokine outputs during DC–T cell priming in an antigen-dependent manner (34), with intracellular NP and secreted sGC antigens exhibiting distinct UTR-sensitivity landscapes and mixed helper polarization profiles when paired with specific UTR configurations. In vivo under fixed ARCA capping, UTR choice reprogrammed NP-specific cellular immunity: CCHFV- and ACTA-1–UTR constructs enhanced IL-2 ELISPOT responses relative to T-UTR, while ACTA-1 broadened CD4⁺ cytokine polarization across IFN-γ, IL-2, IL-4, TNF-α, and IL-17A; CD8⁺ cytokine readouts were comparatively similar across UTRs.

Cap structure further acted as a practical “gain control” for immunogenicity. Across antigens, cap identity primarily scaled response magnitude and humoral output without overriding antigen-defined polarization: under AG capping, NP/NPmut induced Th1-biased IFN-γ ELISPOT responses whereas sGC constructs produced IL-4–dominant profiles; in contrast, CleanCap M6 increased overall T cell response magnitude in NP/NPmut while IL-4 remained undetectable in sGCs constructs. Serology tracked strongly with cap structure: ARCA constructs elicited weak antibody responses, AG preserved antigen-dependent titers (NP negative, NPmut low, sGC strong), and CleanCap M6 further increased antibody titers across all antigens while preserving these relative constraints.

Against this platform-optimization backdrop, antigen architecture defined the dominant biological constraints on protection in this acute, rapidly disseminating infection model. Intracellular NP constructs elicited strong Th1-biased cellular responses and helper priming yet failed to confer protection, whereas secreted sGC constructs generated potent humoral immunity and high survival—reinforcing that protection depends on inducing the effector mechanism required to restrict early viral spread (35), here glycoprotein-targeted antibodies, and that conventional T cell magnitude alone is an unreliable proxy for efficacy. Targeted antigen edits further separated immunogenicity from protection: removing a caspase-3 cleavage motif in NP increased Tfh and GC responses and improved antibody generation in some animals (20,21) yet did not improve survival, while GP38 glycan removal enhanced neutralization potency consistent with improved epitope accessibility (36) but modestly reduced survival versus wild-type, pointing to trade-offs between exposure, conformational stability, and in vivo antibody functionality.

Challenge studies in a stringent immunosuppressed model validated these platform principles end-to-end. Under CleanCap M6 conditions, sGCs alone conferred robust protection (90% survival) with minimal clinical signs, whereas NP and NPmut conferred no protection; combination regimens yielded intermediate, non-additive protection (30–70%), consistent with antigenic competition and shifted immunodominance in multivalent contexts (37). Protective efficacy aligned most closely with antibody quality and GC recruitment rather than with overall T cell magnitude.

Several limitations frame translational scope. DC-based assays are mechanistically informative but cannot fully recapitulate tissue microenvironments or systemic cytokine dynamics in vivo. The immunosuppressed challenge model emphasizes early antibody-mediated control and may underrepresent contexts where cellular immunity contributes more directly. Finally, the dissociation between neutralization potency and survival highlights the need for richer antibody-quality metrics (epitope targeting, avidity, Fc-mediated breadth) and for future studies integrating systems-level profiling, alternative challenge models, and dose-ranging safety assessments.

Taken together, this work positions mRNA–LNP vaccination as a platform optimization problem: lipid chemistry sets innate/inflammatory context, RNA regulatory elements tune magnitude and balance, and antigen properties establish the protective ceiling. By coupling modular control over these axes, we define design principles that extend beyond CCHFV to other emerging pathogens and provide a transferable foundation for rational, rapid, and safer mRNA–LNP vaccine development in future outbreak settings.

## Methods

### mRNA–LNP Design and Experimental Overview

This study systematically evaluated the immunogenicity and protection efficiency of mRNA–LNP formulations targeting CCHFV antigens, focusing on UTRs, cap structures, and ionizable lipids (Figure 1). The role of ionizable lipid chemistry in mRNA delivery and immune activation was also assessed. LNPs were formulated with a first reported used ionizable lipid, BP-104, and compared in vitro to two FDA-approved lipids, SM-102 (Moderna) and ALC-0315 (Pfizer/BioNTech), all encapsulating NPmut mRNA, prior to in vivo evaluation in C57BL/6J mice. Experiments were conducted in a stepwise manner. In the first phase, mRNA constructs encoding the CCHFV nucleocapsid (NP) protein were generated with three distinct 5′/3′ UTR pairs to assess effects on translation, all capped using ARCA (Cap0). In the second phase, mRNAs encoding wild-type NP, a mutant nucleocapsid (NPmut), sGCs, and sGCs with deglycosylated GP38 named sGCs(GP38Δglyc) were synthesized with AG CleanCap (Cap1) and CCHFV-derived UTRs. In the third phase, these same antigens were transcribed using CleanCap M6 (Cap1) to evaluate the impact of additional cap methylation on translation, immunogenicity and protection in the challenge experiment.

### Cell Lines and Culture Conditions

HEK293T (human embryonic kidney; ATCC® CRL-11268™), C2C12 (mouse myoblast; ATCC® CRL-1772™), HepG2 (human hepatocellular carcinoma; ATCC® HB-8065™), and ExpiCHO-S™ (Thermo Fisher Scientific, Cat# A29127) were used to assess mRNA–LNP cytotoxicity, expression, and protein production. HEK293T, C2C12, and HepG2 cells were cultured in DMEM or MEM supplemented with 10% FBS and 1% penicillin-streptomycin at 37 °C and 5% CO₂. ExpiCHO-S™ cells were maintained in ExpiCHO™ Expression Medium in shaking flasks at 37 °C, 8% CO₂, 130 rpm, and passaged every 3–4 days for recombinant GP38 protein production. HEK-Blue™ IFN-α/β (InvivoGen, Cat# hkb-ifnabv2), HEK-Blue™ IL-6 (InvivoGen, Cat# hkb-hil6), and THP1-Dual™ (InvivoGen, Cat# thpd-nfis) were employed to assess type I interferon activity and innate immune activation in response to NPmut mRNA–LNPs formulated with BP-104, SM-102, or ALC-0315. HEK-Blue™ cells were cultured in DMEM with 10% heat-inactivated FBS and antibiotics (penicillin, streptomycin, Normocin®, Blasticidin, Zeocin®) at 37 °C, 5% CO₂. THP1-Dual™ cells were maintained in RPMI 1640 with 10% heat-inactivated FBS, L-glutamine, and antibiotics under the same conditions, passaged every 2–3 days to maintain 2 × 10⁵–1 × 10⁶ cells/mL. DC2.4 murine dendritic cells (Millipore Sigma, Cat#SCC142M) are an immortalized antigen-presenting cell line. Cells were maintained on standard tissue-culture–treated plastic in “DC2.4 expansion medium” consisting of RPMI-1640 supplemented with 10% fetal bovine serum, 1× L-glutamine, 1× non-essential amino acids, 1× HEPES buffer solution, and 0.0054× β-mercaptoethanol, and incubated at 37 °C in a humidified 5% CO₂ atmosphere. All cell lines were routinely tested for mycoplasma and used within 15 passages from authenticated stocks.

### Plasmid Construction and mRNA Templates

Human codon-optimized sequences for NP and sGCs were synthesized (GenScript) in pUC57 plasmids and cloned into pCDNA3.1 via Gibson Assembly to generate pGSCD-NP and pGSCD-Mucin-GP38. Modified constructs included NPmut (DEVD → AEVA) and GP38Δglyc (N-X-S/T → Q-X-S/T) to remove two N-linked glycosylation sites, generated using NetNGlyc (https://services.healthtech.dtu.dk/services/NetNGlyc-1.0/), while the mucin like domain remained unaltered. For protein expression, pGSCD-Mucin-GP38 incorporated C-terminal 6×His and Strep Tag II. For bacterial purification, the codon-optimized NP sequence was cloned into pET-30a (+) to generate pET30-GSCD-NP. A pUC57-PolyA plasmid containing a segmented poly(A) tail (38) was used to introduce T7 promoter, 5′ and 3′ UTRs, Kozak sequence, antigen coding regions, and three stop codons, generating linear DNA templates for in vitro mRNA synthesis.

### Recombinant Protein Production and Purification

Nucleocapsid protein was expressed in Rosetta bacteria transformed with pET30-NP. A single colony was expanded in 20 mL TB with 25 µg/mL kanamycin overnight at 37 °C, then diluted into 1 L TB and grown to OD₆₀₀ ≈ 0.8. Protein expression was induced with 1 mM IPTG for 3 hours at 37 °C. Bacterial pellets were lysed on ice in 20 mM sodium phosphate, 500 mM NaCl, 10 mM imidazole, 1% Triton-X, lysozyme (1 mg/mL), and protease inhibitors, followed by sonication and centrifugation at 23,000 × g for 20 min at 4 °C. Lysates were batch-bound to Ni-NTA agarose for 2 hours at 4 °C, eluted with 250 mM imidazole, buffer-exchanged using 50 kDa Amicon filters, and further purified via size-exclusion chromatography (SEC: Superdex 75 10/300 GL, ÄKTA pure™ 25). Fractions were analyzed by SDS-PAGE, Coomassie-stained, snap-frozen, and stored at –80 °C.

GP38 protein was expressed in ExpiCHO-S™ cells. Cells were passaged three times, seeded at 1 × 10⁶/mL in FectoCHO® CD Expression Medium with 10 mM L-glutamine, and transfected with pGSCD-Mucin-GP38 and Furin plasmids (FURIN (NM_002569) Human Untagged Clone, Origene, Cat number: SC118550) (4:1 ratio) using FectoPro®. Supernatants were harvested when viability dropped below 70%, treated with BioLock to remove biotin, centrifuged, filtered (0.22 µm), and adjusted with Buffer 10x W (1:10) for pH adjustment. Proteins were purified using Strep-Tactin®XT 4Flow® resin and eluted with Buffer BXT, followed by SEC as described for the nucleocapsid.

### mRNA Synthesis

For the first mouse study, ARCA-capped mRNAs encoding wild-type nucleocapsid with three UTR combinations (CCHFV, T, ACTA-1) were synthesized using the HyperScribe™ Co-transcription mRNA Synthesis Kit Plus (ApexBio, Cat# K1407). Plasmid templates were linearized with BstBI at 65 °C for 1 hour, purified (DNA Clean & Concentrator-25, Zymo Research, Cat# D4034), and 1 µg was used in a 25 µL transcription reaction incorporating ARCA, 5mCTP, and ψUTP. For the second experiment, AG-capped mRNAs encoding NP, NPmut, sGCs, and sGCs (GP38Δglyc) were synthesized using mMESSAGE mMACHINE™ T7 with CleanCap™ AG (ThermoFisher, Cat# A57620), substituting TheraPure™ GMP N1-Methylpseudo-UTP. For the third experiment, CleanCap M6-capped mRNAs were generated in 10× reaction buffer (Tris-HCl pH 7.5 400 mM, HCl 150 mM, MgCl₂ 160 mM, DTT 100 mM, Spermidine 21.2 mM) using TriLink CleanCap® M6 (Cat# N-7453-10) and CleanScribe™ RNA Polymerase (TriLink, Cat# E-0107-08), incorporating TheraPure™ GMP N1-Methylpseudo-UTP and 5-Methylcytidine-5’-Triphosphate (TriLink, Cat# N-1014-10). For all caps, Double-stranded RNA was removed via cellulose-based purification, followed by further cleanup using Monarch® Spin RNA Cleanup Kit (NEB, Cat# T2050L), and mRNAs were stored at –80 °C. mRNA integrity was confirmed on 1% low-melting agarose gels in 1× TAE with SYBR Safe dye. Samples (1 µg) were mixed 1:1 with 2× RNA loading dye, heated at 70 °C for 5 minutes, run at 120 V for 45 min, and visualized using a ChemiDoc MP (Bio-Rad).

NP and sGCs mRNAs with a library of UTRs (RNAV8 Bio) for DC-T co-culture experiment were generated by T7 in vitro transcription using a custom NTP mixture (ATP, GTP, CTP and N1-methylpseudouridine triphosphate). The UTR design strategies employed the evaluation of secondary structural elements, presence or absence of RNA binding protein (RBP) sequence motifs, as well as microRNAs. Systematic approaches were employed to identify sequence to function relationships. Reactions contained linearized DNA template, transcription buffer (Tris–HCl or HEPES, pH 8.0), DTT, spermidine, NTPs, T7 RNA polymerase, inorganic pyrophosphatase and RNase inhibitor, and were incubated at 37 °C for 2 h. Residual DNA template was removed with RNase-free DNase I (1 U per 1 μg input DNA; 15–30 min at 37 °C, 300 rpm). mRNA was purified by LiCl precipitation (Ambion AM9480; 2.5 M final LiCl; −20 °C for ≥30 min or overnight), pelleted (≥20,000 × g, 30–60 min), washed three times with ice-cold 70% ethanol, air-dried (5–15 min) and resuspended in nuclease-free water. RNA was quantified by NanoDrop and sized/integrity-checked by Agilent 5200 Fragment Analyzer. Cap 1 was installed enzymatically using Vaccinia Capping System (NEB M2080) and 2′-O-methyltransferase (NEB M0366) per manufacturer instructions. Briefly, 300 μg IVT mRNA was heat-denatured (65 °C, 5 min), snap-cooled on ice, then incubated with capping buffer, GTP, S-adenosylmethionine, RNase inhibitor, Vaccinia capping enzyme and 2′-O-methyltransferase (37 °C, 30 min). Reactions were quenched with EDTA (5 mM final), LiCl precipitated as above, further purified using MEGACLEAR™ (Ambion), and re-quantified/verified by NanoDrop and Fragment Analyzer.

### mRNA Expression Analysis

Expression of M6-capped mRNAs was assessed in HEK293T, C2C12, and HepG2 cells by western blot. Cells were seeded in 6-well plates in complete media one day prior. On the day of transfection, media was replaced with 1 mL Opti-MEM 1 hour before transfection using TransIT®-mRNA Transfection Kit (Mirus Bio, Cat# MIR 2225) according to the manufacturer’s instructions. Cells and supernatants were collected 18 hours post-transfection. For western blot, nucleocapsid-containing lysates and GP38-containing supernatants were mixed 1:1 with 2× Laemmli buffer, boiled at 90 °C for 5 minutes, and loaded on 4–20% Mini-PROTEAN® TGX™ gels. Proteins were transferred to nitrocellulose membranes and blocked with 5% milk in 0.1% PBS-Tween for 30 minutes at room temperature. Membranes were incubated with primary antibodies for 1 hour at room temperature: anti-CCHFV nucleoprotein (1:3000, Abcam, Cat# ab190657) or anti-CCHFV pre-Gn glycoprotein, Clone 13G8 (1:2500, BEI, Cat# NR-40294). Following three washes (5 minutes each) with 0.1% PBS-Tween, membranes were incubated for 1 hour with secondary antibodies: anti-mouse IgG-HRP (Promega, Cat# W4021) for NP or goat anti-rabbit IgG-HRP (Abcam, Cat# ab6721) for GP38. Bands were visualized using SuperSignal™ West Femto Maximum Sensitivity Substrate (ThermoFisher, Cat# 34094).

### mRNA–LNP Formulation and Characterization

Purified mRNAs were encapsulated using Flex-M microfluidics (PreciGenome). The aqueous phase contained 478.1 µg mRNA in 3 mL 100 mM sodium citrate (pH 5.1) and was mixed with 1 mL ethanol phase containing DSPC (BROADPHARM, Cat# BP-25623), cholesterol (Sigma, Cat# C8667), BP-104 (BROADPHARM, Cat# BP-26362), and DMG-PEG2000 (BROADPHARM, Cat# BP-25496) at a molar ratio of 10:38.5:50:1.5. Flow rate ratio was 3:1 with an N/P ratio of 6. Collected LNPs were immediately transferred to 50 mL ice-cold 1× PBS, concentrated, buffer-exchanged using 50 kDa Amicon tubes, filtered through 0.2 µm PVDF Titan syringe filters (Thermo Fisher, Cat# 42204-PV), and stored at –80 °C. LNP hydrodynamic diameter, zeta potential, and polydispersity index (PDI) were determined by dynamic light scattering (DLS) using a Zetasizer Pro Red (Malvern Panalytical, UK). Samples were diluted 10 µL in 1 mL 1× PBS and measured in triplicate; results are reported as mean ± SD. Encapsulation efficiency was quantified using the Quant-iT™ RiboGreen RNA Assay (Thermo Fisher, Cat# R11490) in black 96-well plates (Perkin Elmer, CulturPlate-96 F) (39) and measured on a SpectraMax iD3 (excitation 485 nm, emission 528 nm). The apparent pKa of LNPs was assessed using the LipidLaunch™ LNP Apparent pKa Assay Kit (TNS method; Cayman Chemical, Cat# 702680). LNPs (500 nM) were incubated with 6 µM TNS in 150 µL buffered solutions spanning pH 3–10 (20 mM boric acid, 10 mM imidazole, 10 mM sodium acetate, 10 mM glycylglycine, 25 mM NaCl). Fluorescence was measured on a SpectraMax iD3 (excitation 321 nm, emission 445 nm), and pH was recorded for each well.

### In Vitro Dendritic–T Cells Antigen Presentation Assay

To compare antigen presentation potential of dendritic cells (DCs) using BP-104 LNPs versus FDA-approved ionizable lipids SM-102 (BROADPHARM, Cat# BP-25499) and ALC-0315 (BROADPHARM, Cat# BP-25498), splenic CD11c⁺MHCII⁺ DCs were isolated from naïve 8–10-week-old female C57BL/6J mice using EasySep™ Mouse CD11c Positive Selection Kit II (STEMCELL, Cat# 18781). Cells were resuspended in RPMI supplemented with 10% FBS, 1% penicillin-streptomycin, 0.1 mM β-mercaptoethanol, and 2 mM L-glutamine, plated at 5×10⁵ cells/well in 6-well plates, and stimulated with 5 µg of NPmut mRNA/LNPs formulated with BP-104, SM-102, or ALC-0315. Lipopolysaccharide (100 ng/mL; Stemcell, Cat# 100-1270) served as a positive control. After 24 h, DCs were washed twice with 1x PBS and 1.5×10³ cells were plated per well in 96-well plates. Naïve CD4+ (CD4+CD44lowCD62Lhigh) and CD8+ (CD3+CD8+CD44-CD62L+) T cells were isolated from C57BL/6J spleens using EasySep™ Mouse Naïve CD4+ T Cell Isolation Kit (STEMCELL, Cat# 19765) and EasySep™ Mouse Naïve CD8+ T Cell Isolation Kit (STEMCELL, Cat# 18000), respectively and added to the antigen-pulsed DCs at a 1:1 ratio (1.5×10⁵ cells/well in 200 µL). Co-cultures were incubated for 4 days at 37 °C, followed by treatment with Brefeldin A (BioLegend, Cat# 420601, 1:1000) for 5 hours. Cells were collected and stained for intracellular cytokines as described in the flow cytometry analysis section. Supernatants were analyzed for 13 antiviral cytokines/chemokines using the LEGENDplex™ Mouse Anti-Virus Response Panel (13-plex) kit (BioLegend, Cat# 740622). Briefly, supernatants, assay buffer, and standards were incubated with fluorescently encoded capture beads, followed by biotinylated detection antibodies and streptavidin-PE. Beads were washed, resuspended, and acquired on a flow cytometer with 488 nm and 633 nm lasers. At least 300 events per bead population were collected, and data were analyzed using LEGENDplex™ Data Analysis Software.

To evaluate the impact of UTR selection on T cell priming in vitro, we performed a modified dendritic cell–T cell co-culture assay optimized for CD8+/CD4+ readouts. In these experiments, we used the DC2.4 murine dendritic cell line (Sigma, Cat#SCC142) in place of primary DCs. Following mRNA/LNP exposure, antigen-loaded DCs were harvested from culture surfaces using a cell scraper, combined with isolated naive CD4+ or CD8+ T cells, and plated into untreated 96-well plates at a 1:1 DC:T cell ratio. Co-cultures were maintained for the first 3 days without exogenous stimulation to enrich T cells receiving cognate T cell receptors–dependent activation from antigen-loaded DCs. On day 4, the Berefeldin A was added for CD4+ intracellular cytokines reads. For CD8+ cells, co-cultures were supplemented with anti-CD3 (0.5 µg ml−1; BioLegend, Cat#830301) and anti-CD28 (1 µg ml−1; BioLegend, Cat#302902) along with recombinant IL-2, IL-7 and IL-15 (each 10 ng ml−1; BioLegend, Cat#589106, 581906 and 570306, respectively) to support viability and expansion of activated CD8+ T cells on day 4. Brefeldin A was then added, and intracellular cytokine staining was performed 5 h later.

### Animal Immunogenicity Studies

All procedures were approved by the Broad Institute IACUC (protocol #0376-09-23) and conducted under SPF conditions in ventilated Innovive racks with a 12-hour light/dark cycle, temperature 70–72°F, and 30–40% humidity, following *The Guide for the Care and Use of Laboratory Animals*. Female C57BL/6J and BALB/c mice (Jackson Laboratory), 8–12 weeks old, were randomized into experimental groups and immunized IM into the left thigh with 10 µg of mRNA-LNP in 50 µL sterile 1x PBS. BALB/c mice were used only to evaluate ARCA-capped nucleocapsid antibody responses. All immunogenicity studies employed a prime-boost schedule (Day 0 and Day 14), with euthanasia on Day 28 (ARCA, AG) or Day 32 (CleanCap M6). Negative controls received 1x PBS (ARCA, AG) or empty LNPs (iLNP; CleanCap M6). For ARCA-capped constructs, C57BL/6J mice received three nucleocapsid mRNAs with different UTRs (CCHFV, T, ACTA1), while BALB/c mice received NP and NPmut for antibody evaluation. AG- and CleanCap M6–capped vaccines in C57BL/6J mice encoded with NP, NPmut, sGCs, or sGCs (GP38Δglyc). Each 10 µg dose was administered IM, with blood collected 2–3 weeks post-boost via terminal cardiac puncture. Serum was isolated by centrifugation (1,000×g, 15 min, 4°C), aliquoted, and stored at −80°C until analysis.

### Binding Antibody ELISA

Binding antibodies against NP and GP38 were measured using an in-house ELISA. Plates (Immulon 2 HB) were coated overnight at 4°C with 5 µg/mL antigen in 1× PBS (100 µL/well). Plates were washed once with 0.1% Tween-20 in PBS and blocked with 200 µL ChonBlock ELISA Buffer (Chondrex, Cat #9068) for 2 hours at room temperature. After three washes, serially diluted serum samples (2-fold in blocking buffer, 100 µL/well) were added and incubated 2 hours at room temperature. Plates were washed four times, followed by incubation with HRP-conjugated anti-mouse IgG (H+L) (Promega, Cat #W4021, 1:3000 in ChonBlock Detection Buffer, Cat #90681) for 1 hour at room temperature. After four washes, 100 µL TMB substrate was added for 5–15 minutes, and the reaction was stopped with 100 µL stop solution. Absorbance was measured at 450 nm (SpectraMax iD3). End-point titers were defined as the highest serum dilution producing an OD exceeding the mean + 2 SD of negative control wells.

### Neutralization Assay Using Pseudotyped VSV-M

Replication-incompetent pseudotyped vesicular stomatitis virus (VSV) expressing a modified full M segment of CCHFV was generated (27) to assess neutralizing antibodies in sGCs and sGCs (GP38Δglyc) immunized mice. The modified M sequence was synthesized (GenScript) and cloned into pCDNA3.1 (+) to generate pGSCD-VSV-M. HEK293T cells were seeded in 6-well plates a day prior to transfection. On the following day, media was replaced with 1 mL DMEM per well, and cells were transfected with pGSCD-VSV-M and pCAGGS-G-Kan plasmid (Kerafast, Cat #EH1017). Six hours post-transfection, the media was replaced with DMEM containing 10% FBS. After 24 h, G*ΔG-luciferase rVSV (Kerafast, Cat #EH1020-PM) was added for 1 hour at 37°C, followed by washing with 1× PBS and addition of DMEM with 2% FBS. Anti-G VSV antibody (1:1000; Kerafast, Cat #EB0010) was included to neutralize residual VSV-G in the VSV-M wells. Viral supernatants were collected 24 hours later, aliquoted, and stored at −80°C.

For the neutralization assay, serum samples were serially 2-fold diluted starting at 1:80. Fifty microliters of each dilution were mixed with 50 µL of diluted pseudotyped virus and incubated on ice for two hours before adding to Vero cells in 96-well plates. After 1 hours, the mixture was removed, wells were washed with 1× PBS, and 100 µL phenol-red free DMEM with 2% FBS was added. Plates were incubated for 22 h and viral infection was quantified by adding 100 µL Bright-Glo™ substrate (Promega, Cat #E2620) and reading luminescence on a SpectraMax iD3.

### Liver histopathology and cleaved caspase-3 immunohistochemistry

At the study endpoint, the AG-capped immunized mice were euthanized, and their livers were collected for histopathological evaluation. Tissues were immersion-fixed in 4% paraformaldehyde, processed routinely, embedded in paraffin, and sectioned at 4 µm. For morphology, sections were stained with hematoxylin and eosin (H&E) using standard procedures and evaluated for architectural disruption and injury features. For apoptosis assessment, paraffin sections were deparaffinized in xylene and rehydrated through a graded ethanol series, followed by heat-induced epitope retrieval in citrate buffer pH 6.0 (Abcam, #AB93678). Endogenous peroxidase activity was quenched with hydrogen peroxide (Abcam, AB64218) for 10 minutes, and non-specific binding was blocked with Rodent Blocking Reagent (BioCare Medical, RBM961L) for 30 minutes. Sections were incubated with rabbit anti–cleaved caspase-3 (D175) (Cell Signaling Technology, #9664L) for 60 minutes at 1:800, followed by Rabbit HRP-polymer secondary (BioCare, #RMR622L) for 30 minutes and development with 3,3′-diaminobenzidine (DAB) (Epredia, #TA-125-QHDX) for 5 minutes. Slides were counterstained with hematoxylin, dehydrated, and mounted. Bright-field images were acquired using an Olympus BX53 microscope with identical acquisition settings across groups.

### ELISPOT Assay for Antigen-Specific T Cell Responses

Splenocytes were collected 2–3 weeks post-booster from immunized mice to assess antigen-specific T cell responses. For ARCA-capped constructs, IFN-γ and IL-2 responses were measured, while IFN-γ and IL-4 were assessed for AG- and CleanCap M6-capped constructs. Splenocytes were isolated under sterile conditions, resuspended in complete RPMI (10% FBS, 1% penicillin-streptomycin, 0.1 mM β-mercaptoethanol), and plated in 96-well PVDF-backed ELISPOT plates (XEL485, R&D Systems) pre-coated with capture antibodies for IFN-γ, IL-2, or IL-4. Cells were stimulated with 10 µg/mL of SEC-purified NP or GP38 proteins for 24 hours at 37 °C, 5% CO₂. After incubation, plates were washed with ELISPOT wash buffer (R&D Systems, Cat #895308), incubated with biotinylated detection antibodies for 2 hours at room temperature, followed by streptavidin–alkaline phosphatase conjugate (2 hours), and developed using BCIP/NBT substrate (1 hour). Plates were rinsed with distilled water and dried at 37 °C for 30 minutes. Spot-forming units (SFUs) were quantified using an automated ImmunoSpot analyzer (Cellular Technology Ltd) and expressed as SFUs per seeded cell, with background from unstimulated wells subtracted.

### Antigen recall T cell assay

Spleens were harvested and homogenized into single-cell suspensions using a 70 µm cell strainer in complete RPMI 1640 medium containing 10% FBS, 1% penicillin-streptomycin, and 0.1 mM β-mercaptoethanol. Red blood cells were lysed with ACK buffer to obtain clear single-cell preparations. For antigen-specific stimulation, 2 × 10⁶ splenocytes per well were cultured in 96-well plates with 5 µg/mL NP or GP38 at 37 °C with 5% CO₂. Brefeldin A (5 µg/mL, BioLegend, Cat #420601) was added immediately, and incubation continued for 6 hours. DMSO-treated cells served as negative controls, while 50 ng/mL PMA and 1 µg/mL ionomycin served as positive controls.

Cells were washed with 1× PBS and stained with Live/Dead eFluor 506 Fixable Viability Dye (ThermoFisher, Cat #65-0866-14) for 30 minutes at 4 °C. Fc receptors were blocked with BD Pharmingen™ Purified Rat Anti-Mouse CD16/CD32 (Mouse BD Fc Block™, BD, Cat #553142). Surface staining was performed for 30 minutes at 4 °C in the dark using the following antibodies: CD3-PE-Cy5 (BioLegend, Cat #100274), CD4-PE (ThermoFisher, Cat #12-0043-83), CD8a-FITC (BD, Cat #553031), and CD45R/B220-SuperBright 600 (ThermoFisher, Cat #63-0452-82). Cells were fixed and permeabilized using Cytofix/Cytoperm Kit (BD Biosciences, Cat #554714) for 20 minutes at 4 °C, followed by intracellular staining for 20 minutes with: IFN-γ-PE-Cy7 (BioLegend, Cat #163508), IL-2-APC-Cy7 (BD, Cat #560547), IL-4-PE-CF594 (BD, Cat #562450), TNF-α-AlexaFluor 700 (BioLegend, Cat #506338), and IL-17A-PerCP-Cy5.5 (eBioscience, Cat #45-7177-82). After final washes, cells were acquired on a CytoFlex-M flow cytometer equipped with four lasers. Gating strategy was shown in supplementary figure 7b.

### Lymph Node and Dendritic Cell Flow Cytometry

Mouse inguinal lymph nodes were harvested and mechanically dissociated through a 40 µm cell strainer in complete high-glucose DMEM. Cells were centrifuged at 300 × g for 5 minutes, washed twice with 1× PBS, and counted. For germinal center (GC) B and T follicular helper (TfH) cell staining, 2 × 10⁶ cells per sample were incubated with FVS575V Fixable Viable Dye (BD, Cat #565694) for 20 minutes at 4 °C. After washing with FACS buffer (1% FBS in 1× PBS), cells were stained for 30 minutes at 4 °C with antibody cocktails including BD Pharmingen™ Purified Rat Anti-Mouse CD16/CD32 (Mouse BD Fc Block™, BD, Cat #553142), CD19-NovaFluor Red 710 (ThermoFisher, Cat #M004T02R04-A), CD3-APC-eFluor 780 (ThermoFisher, Cat #47-0031-82), CD4-NovaFluor Yellow 690 (ThermoFisher, Cat #M001T02Y05-A), CD95-AlexaFluor 647 (BD, Cat #563647), GL7-PE (BioLegend, Cat #144608), CD185-PE-Vio615 (Miltenyi Biotec, Cat #130-107-656), CD184 (CXCR4)-PE-Cy7 (ThermoFisher, Cat #25-9991-82), CD279 (PD-1)-PerCP-eFluor 710 (ThermoFisher, Cat #46-9985-82), CD86-SuperBright 645 (ThermoFisher, Cat #64-0862-82), CCR6-Brilliant Violet 510 (BD, Cat #747832), and NP/GP38 labeled with Alexa Fluor 405 and 488. NP- and GP38-AF405/AF488 conjugates were prepared using Alexa Fluor™ 488 NHS Ester (ThermoFisher, Cat #A20000) and Alexa Fluor™ 405 NHS Ester (ThermoFisher, Cat #A30100), purified via Zeba Spin Desalting Columns, 7K MWCO (ThermoFisher, Cat #89877), and stored at 4 °C for up to one month. After staining, cells were washed twice with FACS buffer, fixed in 1% paraformaldehyde, and acquired on a CytoFlex-M cytometer. Gating strategy was shown in supplementary figure 7a.

For dendritic cell (DC) analysis, cells were first stained with Fixable Live/Dead Ghost Dye Violet 540 (Cell Signaling, Cat #72086) for 30 minutes at 4 °C in the dark. After washing, cells were incubated with BD Pharmingen™ Purified Rat Anti-Mouse CD16/CD32 (Mouse BD Fc Block™, BD, Cat #553142) and antibodies including CD3-APC-eFluor 780 (ThermoFisher, Cat #47-0032-82), CD19-APC-eFluor 780 (ThermoFisher, Cat #47-0193-82), NK1.1-APC-eFluor 780 (ThermoFisher, Cat #47-5941-82), Ly-6G-APC-eFluor 780 (ThermoFisher, Cat #47-9668-82), CD11c-VioBright B515 (Miltenyi Biotec, Cat #130-129-314), MHCII-PE Vio615 (Miltenyi Biotec, Cat #130-112-393), Siglec-H-APC REAfinity™ (Miltenyi Biotec, Cat #130-112-297), CD317-PE (BioLegend, Cat #127104), and XCR1-VioBright R720 (Miltenyi Biotec, Cat #130-132-757) for 30 minutes at 4 °C. Cells were fixed with 1% paraformaldehyde for 30 minutes at 4 °C before acquisition on the CytoFlex-M.

### In vivo challenge studies

Groups of C57BL/6J mice (n = 10 per group; five females and five males, 8-10 weeks of age; The Jackson Laboratory, Strain: 000664) were vaccinated IM with 10 µg of the designated single-antigen mRNA vaccine (or 10 µg of each mRNA candidate when two were administered in parallel) on day –35 and boosted two weeks later, 21 days prior to virus challenge. Control animals received iLNP alone. All mice were treated with 2.5 mg per mouse of anti-mouse INFAR-1 monoclonal antibody MAR1-5A3 (Leinco Technologies, Inc., Product No.: I-400, Lot no.: 0425L1135) via the intraperitoneal (IP) route prior to subcutaneous (SC) challenge with lethal CCHFV IbAr10200 (Ref #813730; target dose: 100 TCID₅₀; actual dose: 119 TCID₅₀) on day 0. Mice were housed in a climate-controlled laboratory with a 12 hours day/night cycle; provided sterilized commercially available mouse chow and sterile water ad libitum; and group-housed on autoclaved corn cob bedding (Bed-o’Cobs ¼”, Anderson Lab Bedding) with crinkle paper (Enviro-Dry), soft/ultra absorbent bedding (Carefresh) and cotton nestlets in an isolator-caging system (Tecniplast GM500 cages, West Chester, PA, USA) with a HEPA-filtered inlet and exhaust air supply. Mice were evaluated daily for up to 14 days post-challenge to assess clinical signs of disease and assigned a score ranging from 0–10 based on the following criteria: piloerection, hunched posture, hypoactivity, percent weight loss, abnormal respiration, dehydration, and neurological signs (ataxia, paresis). Euthanasia criteria were met when weight loss exceeded 25% from baseline at day 0 and/or the clinical score reached 10. Mice were humanely euthanized via isoflurane exposure followed by cervical dislocation at study completion or when meeting euthanasia criteria.

All CCHFV infections were performed in biosafety level 4 facilities at the Centers for Disease Control and Prevention (Atlanta, GA, USA). All animal procedures were approved by the CDC Institutional Animal Care and Use Committee (IACUC protocol 3342SPEMOUC) and conducted in accordance with the Guide for the Care and Use of Laboratory Animals. The CDC is fully accredited by the AAALAC-International.

### Data Analysis

All data was organized and processed using Microsoft Excel 2023. Statistical analyses were conducted in GraphPad Prism version 10.0.0 (GraphPad Software, Boston, MA, USA, www.graphpad.com). Paired comparisons (e.g., longitudinal measurements within the same animal group) were evaluated using the two-tailed Wilcoxon signed-rank test, while differences among multiple groups were assessed via one-way ANOVA followed by Tukey’s post hoc multiple comparison test. Flow cytometry antibody panels were designed using FluoroFinder (https://fluorofinder.com/), and all plasmids, DNA fragments, and primers were designed in silico using SnapGene (www.snapgene.com). The secondary structures of mRNA untranslated regions were analyzed with QIAGEN CLC Main Workbench 21.0 (QIAGEN, Aarhus, Denmark). Apparent LNP pKa values were calculated by fitting fluorescence data to the Henderson–Hasselbalch equation in Mathematica (Wolfram Research). Results are presented as mean ± standard deviation (SD), and statistical significance was defined as p < 0.05.

## Author contributions

## Acknowledgements

We thank members of the Sabeti Lab and the Viral Special Pathogens Branch at the Centers for Disease Control and Prevention for their essential contributions to experimental design, in vivo studies, and BSL-4 operations.

## Disclaimer

The findings and conclusions in this report are those of the authors and do not necessarily represent the official position of the Broad Institute of MIT and Harvard and Centers for Disease Control and Prevention.

## Conflict of interest

P.C.S. is a co-founder and equity holder in Delve Biosciences, a board member and equity holder in Polaris Genomics, an equity holder and former advisory board member of NextGenJane, a co-founder and equity holder of Lyra Labs, and an equity holder and former member of the Board of Directors of Danaher Corporation. P.C.S no longer holds a financial interest in Sherlock Biosciences. All potential conflicts are managed in accordance with institutional policy. T.F., A.O., and P.S. are co-inventors on International Patent Application No. PCT/US2025/056332 filed by The Broad Institute. RNAV8 Bio has filed patents covering its novel UTRs and related approaches.

## Financial support

The work at Broad Institute of MIT and Harvard was fully supported by a grant (No. 58-3022-2-031) from the U.S. Department of Agriculture. The challenge experiment was supported in part by CDC Emerging Infectious Disease Research Core Funds.

## Supplementary Data

**Supplementary Figure 1.**
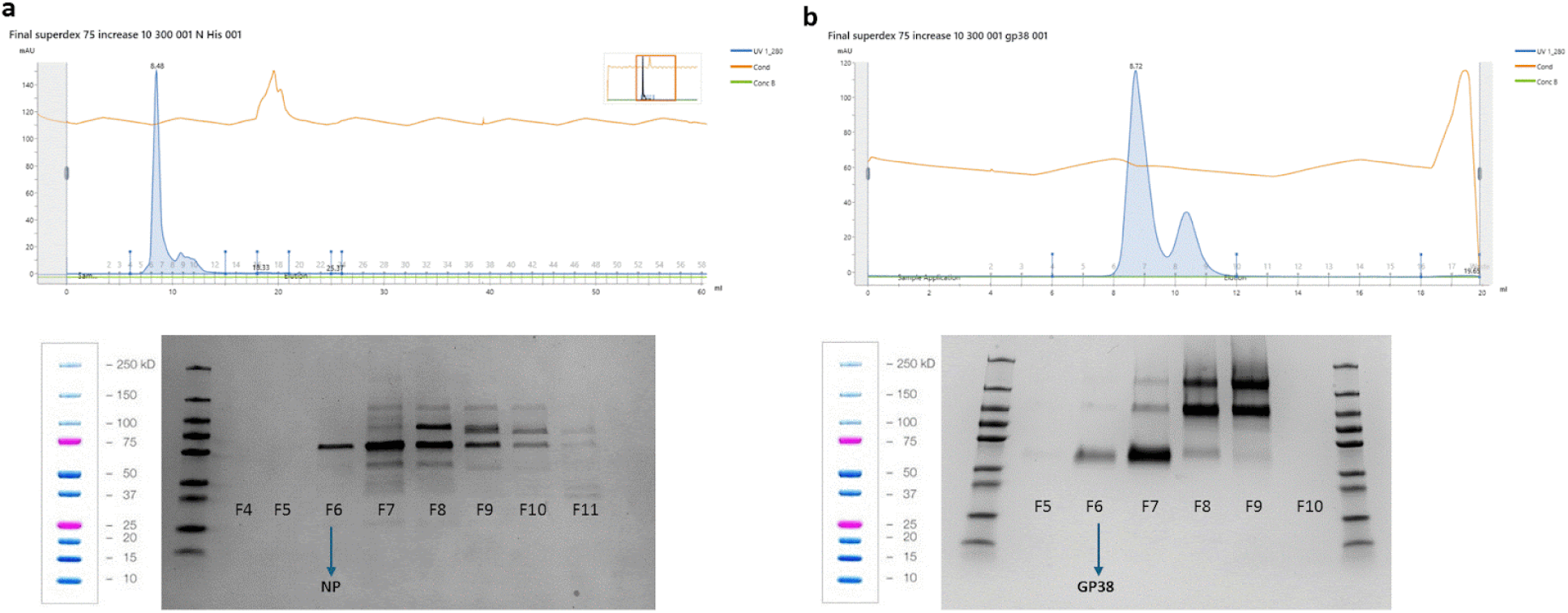
Size exclusion chromatography of recombinant CCHFV proteins. **a,** Nucleocapsid protein was purified from *E. coli* Rosetta cells using Ni-NTA gravity flow and further fractionated by size exclusion chromatography (SEC) on a Superdex™ 75 10/300 GL column (Cytiva). Coomassie blue staining of the collected fractions indicates that peak 6 corresponds to the pure nucleocapsid protein. **b,** Recombinant GP38 was expressed in ExpiCHO cells, purified from culture supernatant using StrepTactin affinity beads, and further separated by SEC on the same Superdex™ 75 column. Coomassie staining shows that peak 6 contains the pure GP38 fraction.

**Supplementary Figure 2.**
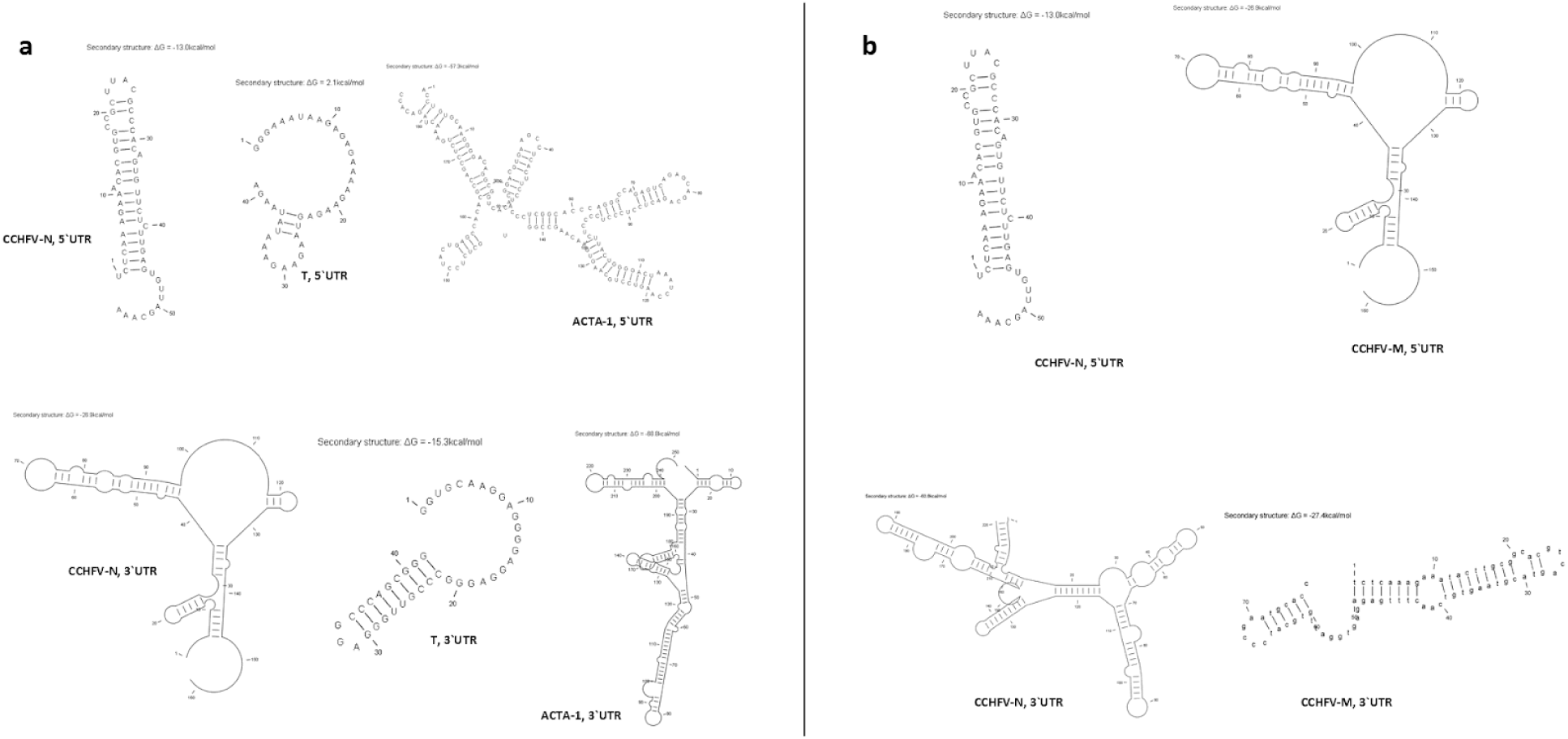
Design and structural analysis of untranslated regions (UTRs) in mRNA vaccine constructs in vivo. **a,** ARCA-capped mRNAs encoding nucleocapsid were engineered with three distinct UTRs: the S-segment UTR from CCHFV (CCHFV-N UTR), an optimized synthetic UTR (T UTR), and the human ACTA-1 gene UTR (ACTA-1 UTR). **b,** Secondary structures of UTRs derived from the S and M segments of CCHFV were incorporated into AG- and M6-capped mRNAs encoding nucleocapsid, mutant nucleocapsid (NPmut), sGCs, and sGCs (GP38Δglyc). The predicted RNA secondary structures, generated using QIAGEN CLC Main Workbench 21.0, illustrate differences in structural complexity among the UTRs, which may influence mRNA stability and translational efficiency.

**Supplementary Figure 3.**
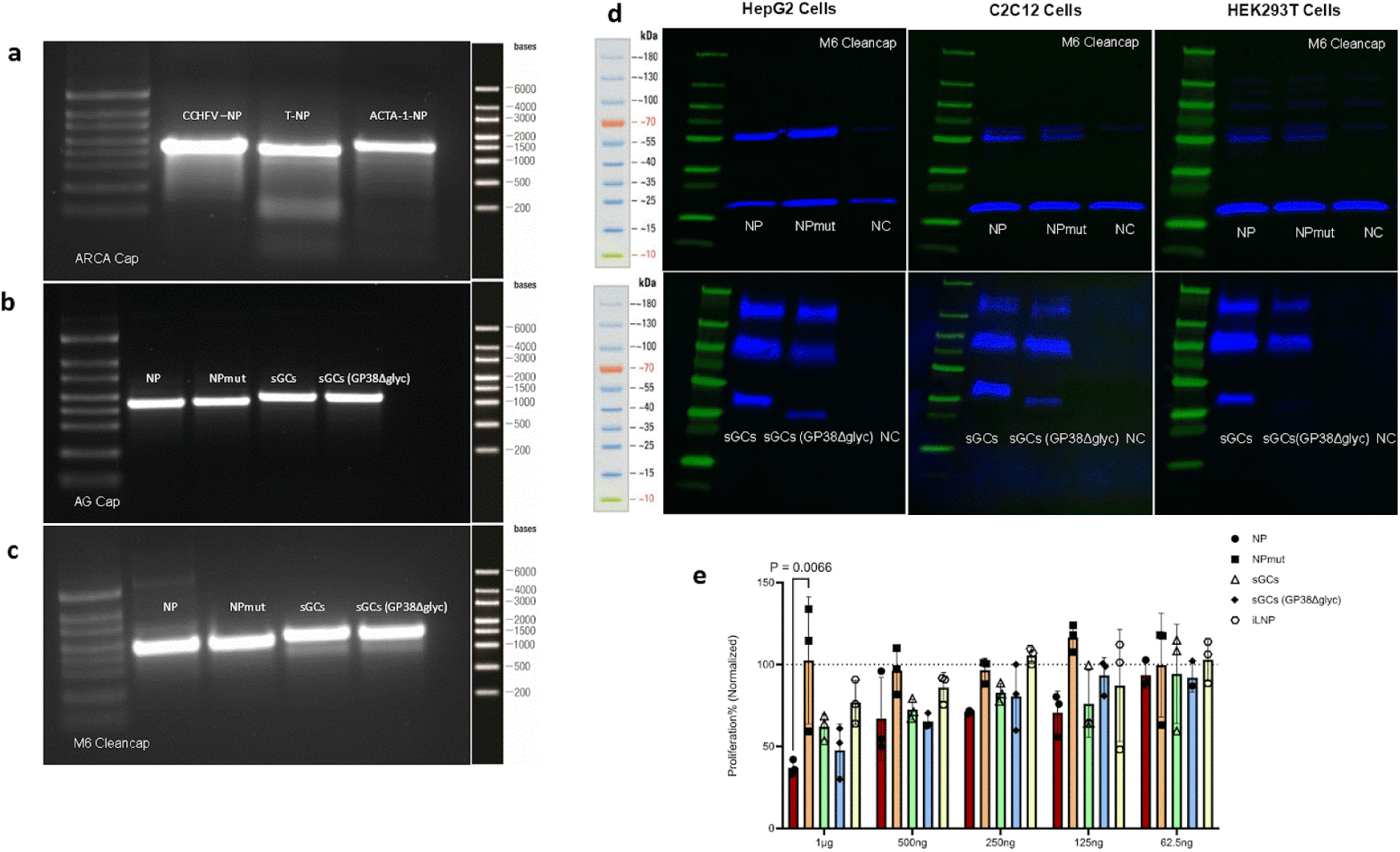
Quality control and in vitro expression of CCHFV mRNA vaccine constructs across cap chemistries and antigen designs. **a–c**, Representative agarose gel electrophoresis of in vitro–transcribed (IVT) mRNAs generated for in vitro and in vivo studies, shown under the three cap-structure conditions used throughout the study. **a,** ARCA-capped (Cap0) nucleocapsid (NP) mRNAs carrying distinct untranslated region (UTR) configurations (CCHFV-UTR–NP, T-UTR–NP, and ACTA-1–UTR–NP). **b,** AG-capped (Cap1) mRNAs encoding NP, NPmut, the secreted glycoprotein complex (sGCs; MLD–GP38), and the sGCs(GP38Δglyc). **c,** CleanCap M6–capped (Cap1) mRNAs encoding the same antigen panel (NP, NPmut, sGCs, and sGCs(GP38Δglyc)). Size ladders (bases) are shown adjacent to each gel. The presence of single dominant bands at the expected transcript sizes indicates intact, full-length IVT products across UTR, antigen, and cap-structure configurations. **d,** Immunoblot analysis of antigen expression following transfection with CleanCap M6 mRNAs in three cell types (HepG2, C2C12, and HEK293T). For nucleocapsid constructs (top row), lysates from cells transfected with NP or NPmut mRNAs are shown alongside a negative control (NC). For glycoprotein constructs (bottom row), supernatants from cells transfected with sGCs or sGCs(GP38Δglyc) mRNAs are shown alongside NC. Molecular weight markers (kDa) are displayed, and bands corresponding to expressed antigens are visible across cell types, indicating robust translation of both intracellular (NP, NPmut) and secreted glycoprotein (sGCs, sGCs(GP38Δglyc)) immunogens under the M6 cap architecture. **e,** In vitro cytotoxicity/tolerability profiling across antigen-encoding mRNA/LNP formulations. Cell viability (normalized proliferation, %) was quantified across a dose range (1 μg to 62.5 ng) for LNPs encapsulating NP, NPmut, sGCs, or sGCs(GP38Δglyc) mRNAs, with empty LNPs (iLNP) included as a formulation control. Individual replicates are overlaid on bar summaries; the dotted line denotes the normalization reference. A modest increase in cytotoxicity was observed for specific antigen–formulation pairings at higher doses (P value indicated), whereas most conditions clustered near the iLNP baseline across the dilution series, supporting broadly favorable in vitro tolerability of the mRNA/LNP preparations.

**Supplementary Figure 4.**
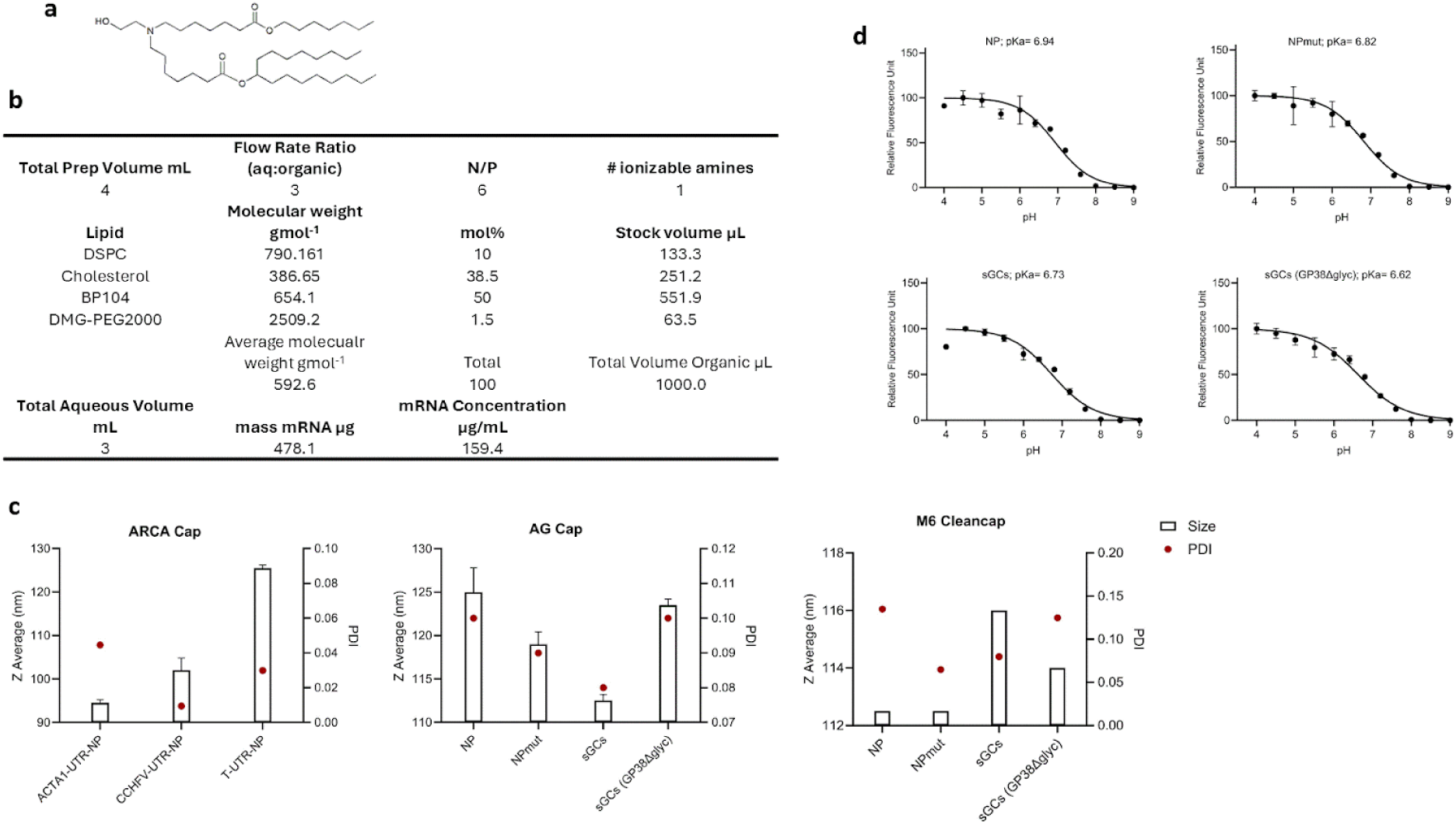
BP-104 lipid composition supports formation of uniform mRNA–LNPs with consistent ionization properties across antigen and cap-structure designs. **a,** Chemical structure of the ionizable lipid BP-104 used for LNP formulation throughout the study. **b,** Representative formulation parameters for BP-104 LNP preparation, including total preparation volume, aqueous:organic flow-rate ratio, N/P ratio, and lipid composition (DSPC, cholesterol, BP-104, and DMG-PEG2000) with molecular weights, mol% and stock volumes used to generate LNPs under standardized mixing conditions. Total aqueous volume, total mRNA mass, and resulting mRNA concentration are indicated. **c,** Dynamic light scattering (DLS) characterization of BP-104 mRNA–LNPs formulated with distinct antigens and cap architectures. Plots show Z-average hydrodynamic diameter (bars; nm) and polydispersity index (PDI; red points) for ARCA-capped NP constructs with alternative UTRs (ACTA-1-UTR–NP, CCHFV-UTR–NP, T-UTR–NP; left), AG-capped antigen panel (NP, NPmut, sGCs, and sGCs(GP38Δglyc); middle), and CleanCap M6–capped antigen panel (right). Across cap structures and antigen configurations, BP-104 LNPs formed monodisperse particles with low PDI values, indicating consistent nanoparticle uniformity. **d,** Apparent pKa measurements for BP-104 mRNA–LNPs encapsulating NP, NPmut, sGCs, or sGCs(GP38Δglyc), determined by fluorescence-based titration across pH. Curves show relative fluorescence as a function of pH with fitted sigmoidal transitions; the derived pKa values are indicated for each construct. BP-104 LNPs exhibited pKa values in the range associated with effective endosomal escape, and these ionization characteristics were maintained across antigen designs.

**Supplementary Figure 5.**
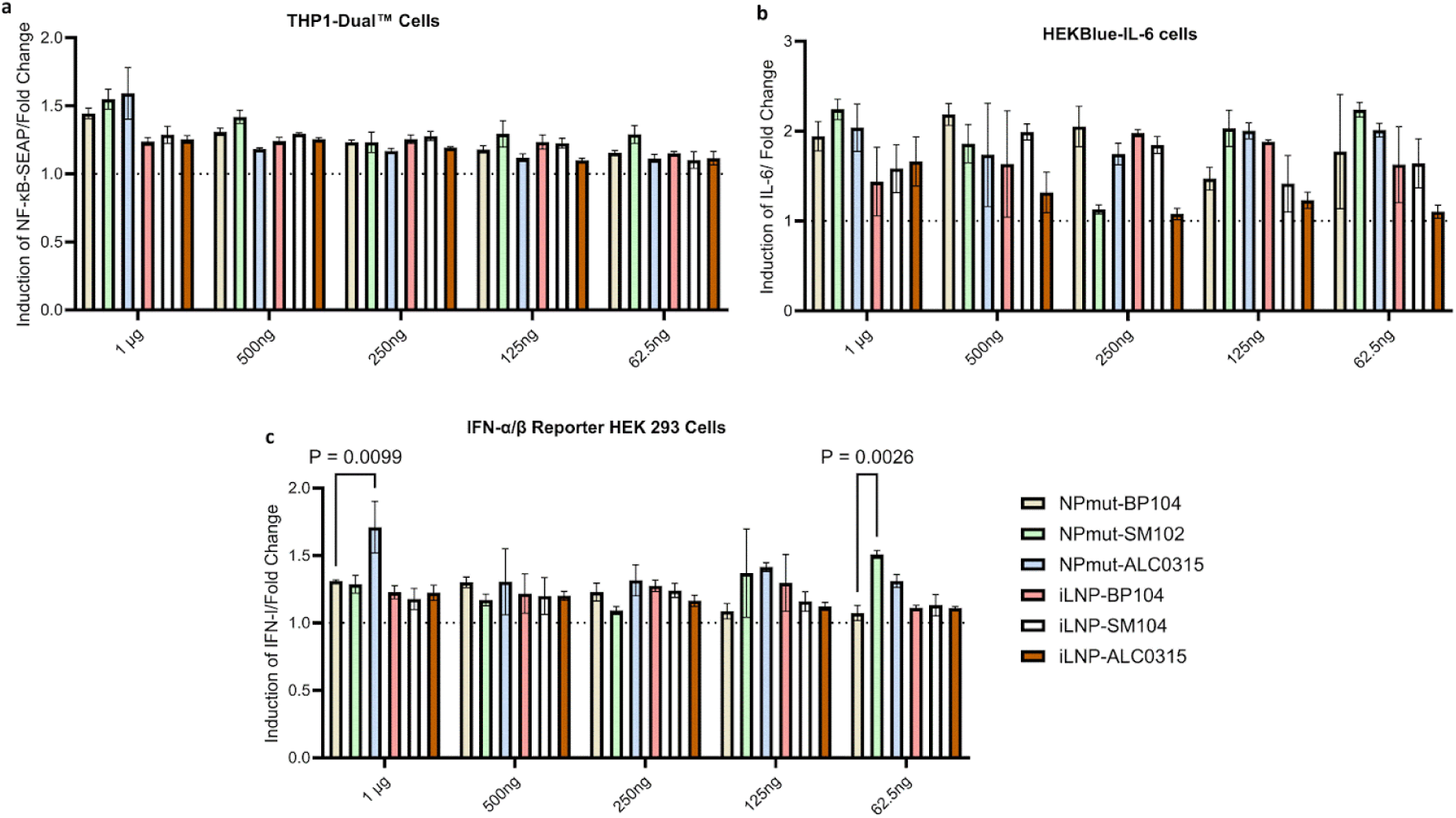
BP-104 elicits innate pathway activation comparable to clinically deployed ionizable lipids across reporter systems. Innate activation was compared across mRNA–LNPs formulated with BP-104, SM-102, or ALC-0315 using NPmut mRNA as a common payload, and across matched empty LNP controls (iLNP; no mRNA) to isolate lipid-driven signaling. Reporter cells were exposed to a dose range of formulations (1 μg, 500 ng, 250 ng, 125 ng, and 62.5 ng; as indicated), and pathway activation was quantified as fold-change induction relative to baseline (dotted line). Bars denote mean responses with error bars indicating variability across replicates; selected pairwise comparisons are annotated with exact P values. **a,** NF-κB activation measured in THP1-Dual™ cells as induction of NF-κB–SEAP. Across doses, NPmut mRNA/LNPs and empty LNPs produced modest but consistent NF-κB induction, with broadly similar activation profiles across BP-104-, SM-102-, and ALC-0315–containing LNPs. **b,** IL-6 pathway signaling measured in HEKBlue–IL-6 reporter cells as fold-change induction of IL-6–dependent signaling. All three ionizable lipid formulations elicited comparable IL-6 activation across the dilution series, and empty LNP controls also induced IL-6 signaling, indicating that both lipid composition and mRNA cargo contribute to pathway engagement. **c,** Type I interferon signaling measured in IFN-α/β reporter HEK293 cells as fold-change induction of IFN-I activity. NPmut mRNA/LNPs triggered IFN-I responses across lipids, with ALC-0315 producing slightly higher induction at select concentrations (P values shown). Empty LNPs also elicited measurable IFN-I signaling, consistent with lipid-dependent innate stimulation independent of mRNA. Collectively, these data show that BP-104 engages canonical innate immune pathways (NF-κB, IL-6, and type I interferon) at levels comparable to SM-102 and ALC-0315 under controlled dosing conditions, supporting its suitability as an ionizable lipid for mRNA–LNP vaccine delivery.

**Supplementary Figure 6.**
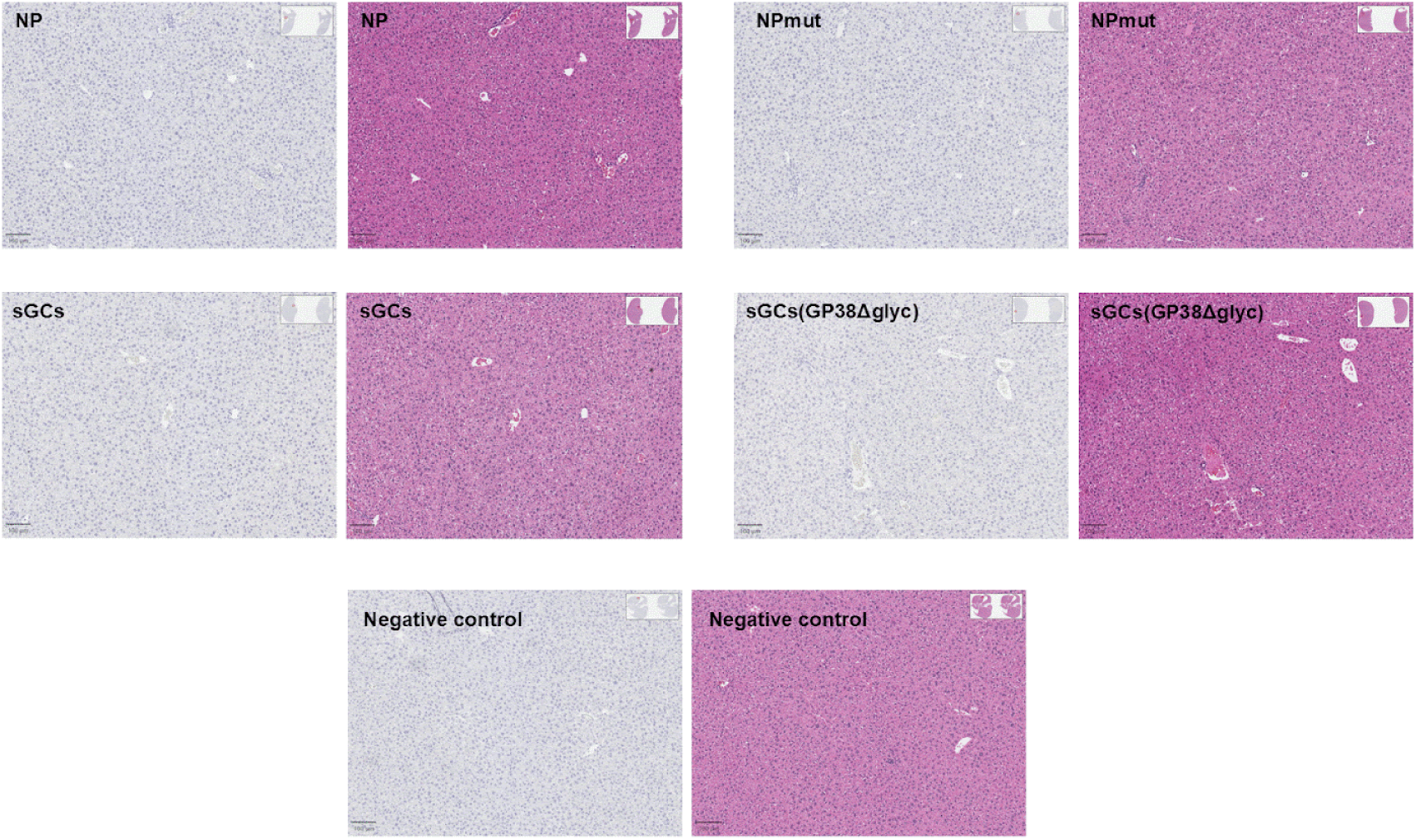
Liver histopathology and apoptosis immunohistochemistry at study endpoint following AG-capped mRNA–LNP immunization. At endpoint, livers from AG-capped–immunized mice (NP, NPmut, sGCs, or sGCs(GP38Δglyc)) and negative-control animals were collected, immersion-fixed in 4% paraformaldehyde, paraffin embedded, and sectioned at 4 μm. Representative H&E-stained sections (right) show hepatic morphology and were evaluated for architectural disruption and injury. Representative immunohistochemistry for apoptosis (left) was performed on deparaffinized sections after heat-induced epitope retrieval in citrate buffer (pH 6.0), endogenous peroxidase quenching, and rodent blocking, using rabbit anti–cleaved caspase-3 (D175; 1:800) with HRP-polymer detection and DAB chromogen (brown), with hematoxylin counterstain (blue nuclei). Bright-field images were acquired on an Olympus BX53 microscope using identical settings across groups. Scale bars, 100 μm.

**Supplementary Figure 7.**
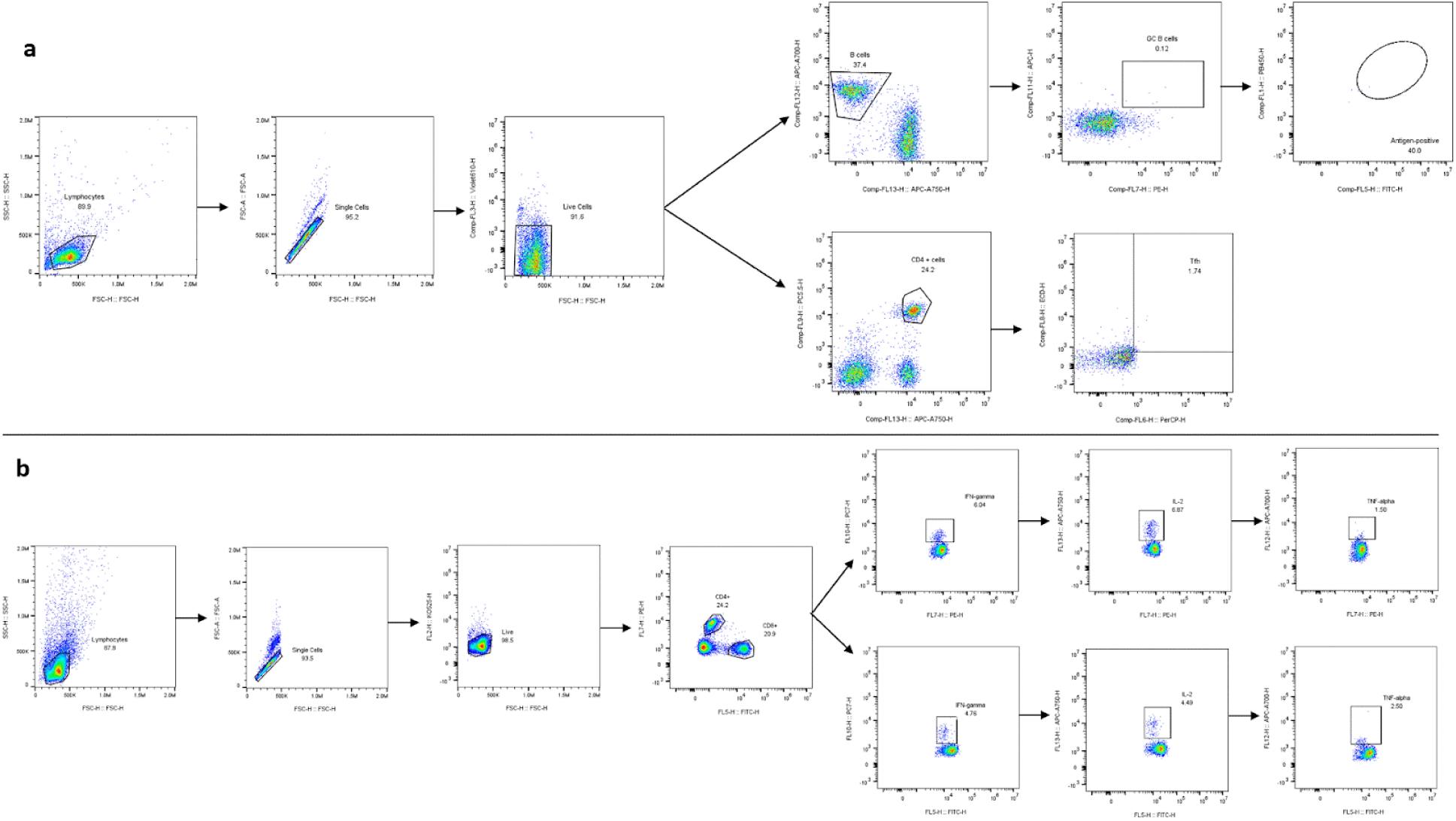
Flow cytometry gating strategies for immune cell analysis. **a,** Gating strategy used to identify T follicular helper (Tfh) cells, germinal center (GC) B cells, and antigen-specific GC B cells in inguinal lymph nodes of immunized mice. **b,** Gating strategy applied to splenocytes to quantify intracellular cytokine production in CD4⁺ and CD8⁺ T cells following ex vivo stimulation.

## Notes

### Competing Interest Statement

The authors have declared no competing interest.

